# Integrative analysis of the plasma proteome and polygenic risk of cardiometabolic diseases

**DOI:** 10.1101/2019.12.14.876474

**Authors:** Scott C. Ritchie, Samuel A. Lambert, Matthew Arnold, Shu Mei Teo, Sol Lim, Petar Scepanovic, Jonathan Marten, Sohail Zahid, Mark Chaffin, Yingying Liu, Gad Abraham, Willem H. Ouwehand, David J. Roberts, Nicholas A. Watkins, Brian G. Drew, Anna C. Calkin, Emanuele Di Angelantonio, Nicole Soranzo, Stephen Burgess, Michael Chapman, Sekar Kathiresan, Amit V. Khera, John Danesh, Adam S. Butterworth, Michael Inouye

## Abstract

Common human diseases are frequently polygenic in architecture, comprising a large number of risk alleles with small effects spread across the genome^1–3^. Polygenic scores (PGSs) aggregate these alleles into a metric which represents an individual’s genetic predisposition to a specific disease. PGSs have shown promise for early risk prediction^4–7^, and there is potential to use PGSs to understand disease biology in parallel^8^. Here, we investigate the role plasma protein levels play in cardiometabolic disease risk in a cohort of 3,087 healthy individuals using PGSs. We found PGSs for coronary artery disease (CAD), type 2 diabetes (T2D), chronic kidney disease (CKD), and ischaemic stroke (IS) were associated with levels of 49 plasma proteins. These associations were polygenic in architecture, largely independent of *cis* protein QTLs, and robust to environmental variation. Over a median 7.7 years follow-up, 28 of these plasma proteins were associated with future myocardial infarction (MI) or T2D events, 16 of which were causal mediators between polygenic risk and incident disease. These protein mediators of polygenic disease risk included targets of approved therapies which may have repurposing potential. Our results demonstrate that PGSs can identify proteins with causal roles in disease, and may have utility in drug development.

## Main Text

Cardiometabolic diseases have a major polygenic component, which is due to the combination of many thousands of variants across the genome, each exerting small lifelong effects^9–13^. Risk stratification using cardiometabolic PGSs have shown potential clinical utility for disease prevention^14^; however, molecular mediators of polygenic risk and their potential to be modulated to reduce disease risk remains unknown. Variants associated with polygenic traits are spread across many different pathways, exerting their effects through multiple levels of regulation, including gene expression, proteins and their interactions, cell morphology, and higher order physiological processes^15^. Proteins that are pathway-level hubs through which polygenic effects converge, however, could be promising targets for pharmaceutical intervention^16– 19^.

Here, we demonstrate how PGSs can be used to identify proteins with causal roles in disease aetiology. The INTERVAL cohort comprises approximately 50,000 adult blood donors in England^20,21^, of which 3,087 participants have linked electronic hospital records, imputed genome-wide genotypes, and quantitative levels of 3,438 plasma proteins^22^ (**Online Methods, Supplementary Data 1**,**2**). The characteristics of the participants are given in **Extended Data Table 1**; and participants with history of any cardiometabolic disease were excluded (**Online Methods, Supplementary Table 1**), reducing the potential for reverse causality in downstream analysis. A schematic of the study is given in **Extended Data Fig. 1**.

To quantify each participant’s relative polygenic risk of atrial fibrillation (AF), CAD, CKD, IS, and T2D we applied externally derived genome-wide PGSs comprised of 1.8–3.2 million variants (**Online Methods**). Using PGSs, we identified 49 proteins whose levels differed with respect to polygenic risk at a false discovery rate (FDR) of 5% (**Fig. 1a,b, Extended Data Table 2,3, Supplementary Table 2**,**3**): 31 proteins for the T2D PGS, 11 proteins for the CAD PGS, 1 protein for the IS PGS, and 8 proteins for the CKD PGS. Associations included proteins with established roles in cardiometabolic disease, such as cystatin-c (CST3) and beta-2-macroglobulin (B2M) which are biomarkers for chronic kidney disease^23^, apolipoprotein E (APOE) whose link to coronary artery disease has been extensively studied^24,25^, and fructose-1,6-bisphosphatase 1 (FBP1) which plays a key role in glucose regulation and is a target of type 2 diabetes drugs^26^. Associated proteins belonged to multiple non-overlapping pathways (**Supplementary Information**), and many are relatively understudied in the context of their respective diseases (**Extended Data Table 4**) warranting future study.

**Figure 1:**
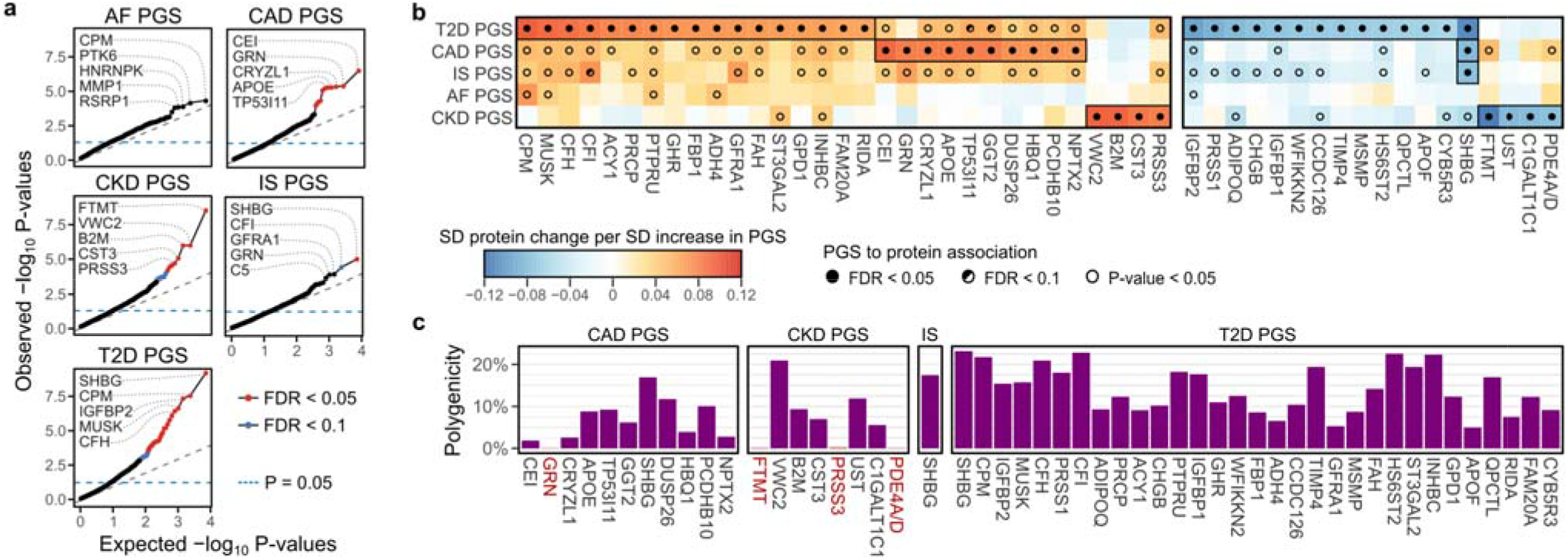
Proteins associated with polygenic risk for cardiometabolic disease. **a)** Quantile-quantile plots of P-values for PGS to protein associations across all 3,438 tested proteins. Each plot compares the distribution of observed P-values (y-axes) to the distribution of expected P-values under the null-hypothesis for 3,438 tests (x-axes) on a –log_10_ scale. Associations were adjusted for age, sex, 10 genotype PCs, sample measurement batch, and time between blood draw and sample processing. Full summary statistics are provided in **Supplementary Data 3a. b)** Heatmaps showing the 49 proteins whose levels significantly associated (FDR < 0.05) with at least one PGS. Each heatmap cell shows the standard deviation change in protein levels per standard deviation increase in PGS, estimated linear regression adjusted for age, sex, 10 genotype PCs, sample measurement batch, and time between blood draw and sample processing. Proteins are ordered by PGS from left to right by decreasing association magnitude, positive and negative associations split into separate heatmaps. Point estimates are detailed in **Extended Data Table 2**. Details about each protein are provided in **Extended Data Table 3. c)** Barplots showing the proportion of the genome required to explain each PGS to protein association (polygenicity; **Online Methods**). Proteins are ordered from left to right by strength of PGS to protein association. Highlighted in red are PGS to protein associations that were explained by singular variants regulating the protein levels, protein quantitative trait loci (pQTLs), rather than polygenic.

PGS to protein associations were robust to technical, physiological, and environmental confounding (**Supplementary Information**). We observed directional consistency and strong correlation of effect sizes when utilizing an orthogonal proteomics technology in independent samples (**Extended Data Fig. 2a-c**). Protein levels and PGS to protein associations were also temporally stable over two years of follow-up (**Extended Data Fig. 2c-d**). PGS to protein associations were also robust to circadian and seasonal effects, inclusion of participants with any prevalent cardiometabolic disease, and body mass index (BMI), with the exception of six T2D PGS to protein associations that were partially mediated by BMI (**Extended Data Fig. 2f-g**).

Most PGS to protein associations were not explained by protein quantitative trait loci (pQTLs) but instead were highly polygenic (**Online Methods**; **Fig. 1c**): each protein required a median 12% of the genome to explain its association with a PGS (**Fig. 1c**). Only four associations could be explained by pQTLs, and contributing loci were spread across the genome for the remaining 46 (**Extended Data Fig. 3**). Interestingly, the effects of PGSs and pQTLs on protein levels were largely independent (**Online Methods, Supplementary Table 4**), suggesting that polygenic risk can enhance or buffer locus-specific effects on protein levels.

Three possible scenarios could explain a PGS to protein association^27^: (1) the protein plays a causal role in disease, (2) the protein levels are changing in response to disease processes, but are not themselves causal (reverse causality), and (3) the protein levels are correlated with some other causal factor (confounding) (**Fig. 2a**). Utilizing a median of 7.7 years of follow-up in nation-wide electronic hospital records, we examined whether levels of PGS-associated proteins were associated with risk of onset of the respective cardiometabolic disease, then performed mediation analysis^28^ to identify the proteins that mediate PGS to disease associations, and thereby play causal roles in disease pathogenesis (**Online Methods**). Limited by the number of incident disease events, we restricted our analyses to CAD and T2D (**Extended Data Fig. 4**). 25 of 31 (81%) of T2D PGS proteins were significantly associated (P < 0.05) with increased risk of T2D and 3 of 11 (27%) of CAD PGS proteins were significantly associated with increased risk of incident MI (**Fig. 2b, Extended Data Table 2**). There was directional consistency and strong correlation (Pearson correlation: 0.96, P = 4×10^−23^) between effects of PGSs on protein levels and hazard ratios for protein levels on incident disease risk (**Fig. 2c**). Using mediation analysis, we found that one and 15 proteins were significant mediators between polygenic risk of MI and T2D, respectively, indicating causal roles in disease pathogenesis (**Fig. 2d**).

**Figure 2:**
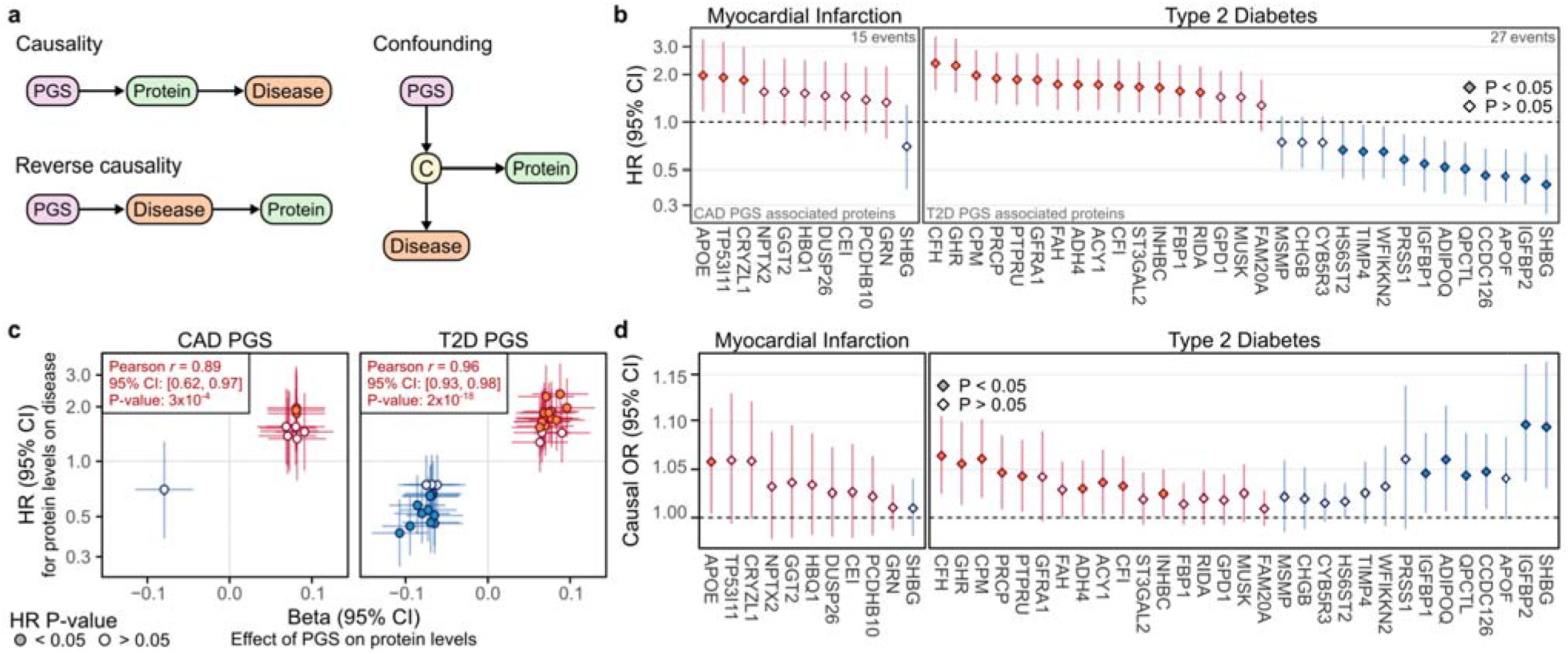
PGS-associated proteins influence 7.7 year risk of myocardial infarction and diabetes. **a)** Possible models of causality for PGS to protein to disease associations. C: causal disease factor upstream of protein that induces a correlation between protein levels and disease. **b)** Association between PGS-associated proteins with 7.7 year risk of hospitalisation with myocardial infarction and diabetes. There were insufficient events to analyse proteins associated with the IS PGS (N=3 incident disease events) or with the CKD PGS (N=0 incident disease events) (**Extended Data Fig. 4a**). Cox proportional hazard models were fit between protein levels and incident disease using follow-up as time scale and adjusting for age and sex (**Online Methods**). HR: hazard ratio conferred per standard deviation increase in protein levels. 95% CI: 95% confidence interval. See **Extended Data Table 2** for detailed point estimates. **c)** Comparison of effects of PGS on protein levels (x-axes; **Fig. 1b**) to associations between protein levels and incident disease (y-axes; **Fig. 2b**). Points and horizontal bars on the x-axes indicate standard deviation change in protein levels (and 95% confidence interval) per standard deviation increase in respective PGS. Points and vertical bars on the y-axis show hazard ratio (and 95% confidence interval) per standard deviation increase in protein levels. **d)** Estimated causal effect of PGS on disease through each protein in mediation analysis (**Online Methods**). Causal OR: odds ratio for incident disease adjusting for age and sex conferred through each protein per standard deviation increase in PGS. The total odds ratio for MI conferred per standard deviation increase in CAD PGS was 2.94 (95% CI: 1.69–5.31, P-value: 2×10^−4^). The total odds ratio for T2D conferred per standard deviation increase T2D PGS was 2.00 (95% CI: 1.37–2.96, P-value: 4×10^−4^). Proteins are ordered from left to right by their hazard ratio in **Fig. 1b. b-d**) points in red indicate proteins whose levels increased with PGS, and blue indicates proteins whose levels decreased with PGS.

As polygenic disease risk is itself estimated from population-level data, it is unlikely that any single protein explains polygenic risk. Here, we found that causal protein mediators each explained a median of 6.6% of PGS to disease associations (**Extended Data Table 2**), with the 1 CAD PGS mediator (APOE) explaining 5.4% of CAD polygenic risk to incident MI association, and the 15 T2D PGS mediators explaining 27% of the T2D polygenic risk to incident T2D association. A complementary approach for causal inference, Mendelian randomisation^29^ (**Online Methods**), also supported causal effects on T2D for two proteins (SHBG and CFI) which mediated the T2D PGS to T2D association (**Supplementary Information, Extended Data Fig. 5, Supplementary Table 5, 6**). Notably, only 12 (24%) of the proteins associated with PGSs could be tested with Mendelian randomisation due to lack of *cis* protein quantitative trait loci (pQTLs) as genetic instruments (**Online Methods**) highlighting the complementarity of our PGS-protein association approach.

Finally, to identify druggable targets associated with polygenic disease risk and potential drug repurposing opportunities, we utilised the DrugBank database^30^ (**Online Methods**) to find that 18 of the 49 PGS-associated proteins were targeted by 236 drugs (**Extended Data Table 5, Supplementary Table 7**). Ten licensed drugs had protein target effects which were consistent with reduction of cardiometabolic disease risk (**Table 1**). These included the well-known T2D drug metformin^31^, which reduces liver glucose production by inhibition of FBP1^32^, a protein whose levels were elevated in people with high polygenic risk for T2D (**Fig. 1**). Among the other nine licensed drugs, we highlight the potential to repurpose pegvisomant for T2D prevention. Pegvisomant (DB00082) is used to treat acromegaly by blocking the binding of endogenous growth hormone to growth hormone receptor (GHR)^33–35^. We found increased GHR was a causal mediator of polygenic T2D risk and incident T2D (**Fig. 2**) and GHR loss-of-function mutations are associated with lower T2D risk^36^ providing additional genetic support for this target. Furthermore, pegvisomant has been shown to improve insulin sensitivity in acromegaly patients^37,38^. Together, these observations suggest pegvisomant is a priority to evaluate for repurposing for T2D prevention (**Table 1**).

**Table 1:**
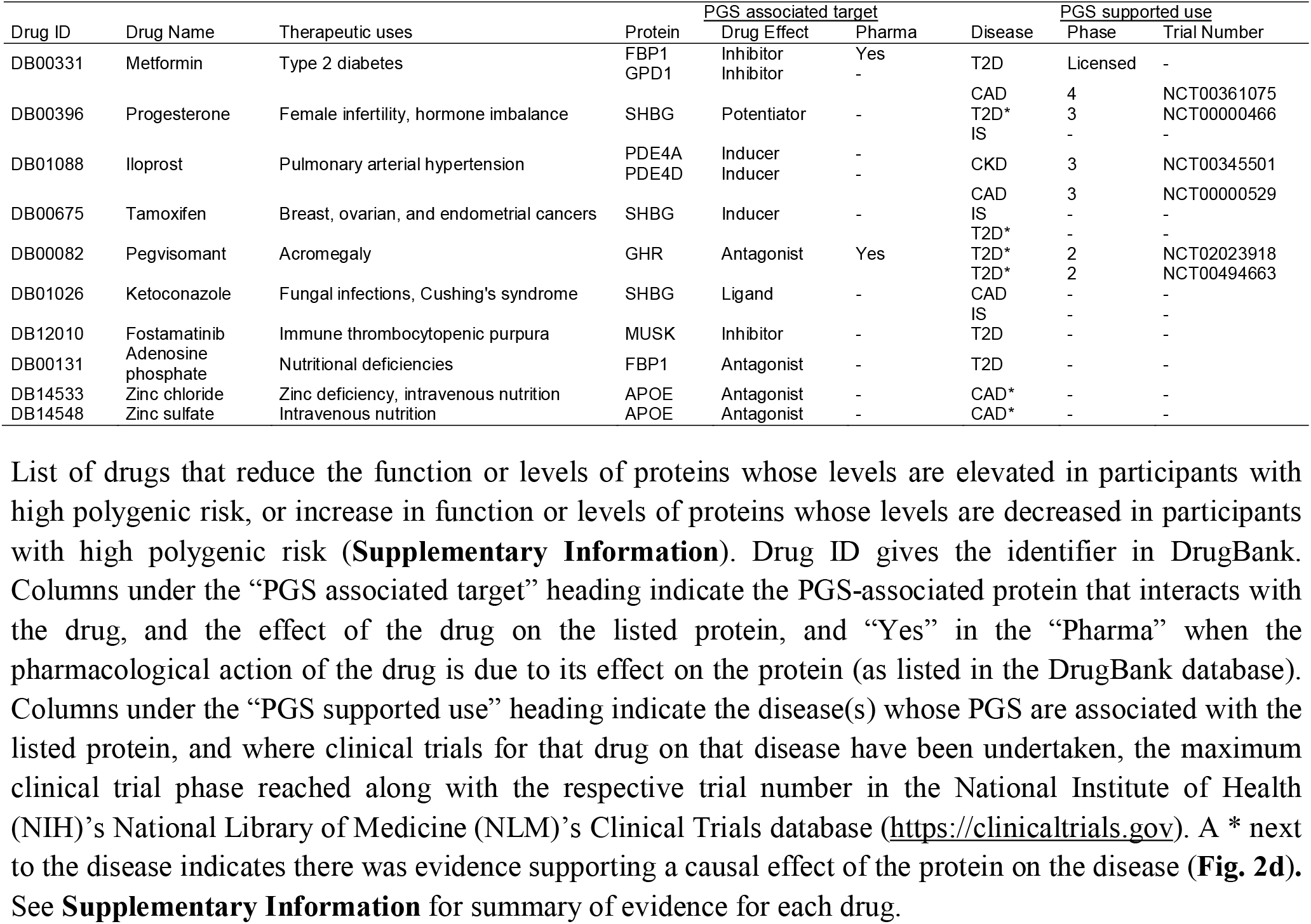
Drugs whose effects on proteins counteract effects of PGSs on proteins. List of drugs that reduce the function or levels of proteins whose levels are elevated in participants with high polygenic risk, or increase in function or levels of proteins whose levels are decreased in participants with high polygenic risk (**Supplementary Information**). Drug ID gives the identifier in DrugBank. Columns under the “PGS associated target” heading indicate the PGS-associated protein that interacts with the drug, and the effect of the drug on the listed protein, and “Yes” in the “Pharma” when the pharmacological action of the drug is due to its effect on the protein (as listed in the DrugBank database). Columns under the “PGS supported use” heading indicate the disease(s) whose PGS are associated with the listed protein, and where clinical trials for that drug on that disease have been undertaken, the maximum clinical trial phase reached along with the respective trial number in the National Institute of Health (NIH)’s National Library of Medicine (NLM)’s Clinical Trials database (https://clinicaltrials.gov). A * next to the disease indicates there was evidence supporting a causal effect of the protein on the disease (**Fig. 2d**). See **Supplementary Information** for summary of evidence for each drug.

## Conclusions

Polygenic scores for disease are explicitly constructed to maximise risk prediction, typically without consideration of the underlying biology. However, PGSs also hold considerable promise for identifying molecular pathways in the development and progression of disease^8,27^. Here, we identified plasma proteins significantly associated with PGSs for cardiometabolic disease in a healthy pre-disease cohort. The vast majority of these associations were highly polygenic, revealing an unappreciated role for polygenic effects on protein levels, including for several well-known disease proteins. These proteins were predictive of incident disease, and 16 were mediators of type 2 diabetes or myocardial infarction, suggesting that their modulation is likely to attenuate disease risk. There are multiple licensed drugs for many of these targets. Overall, this study demonstrates the power of polygenic scores to elucidate novel disease biology and their potential to inform development of medicines.

## Acknowledgements

Participants in the INTERVAL randomised controlled trial were recruited with the active collaboration of NHS Blood and Transplant England (www.nhsbt.nhs.uk), which has supported field work and other elements of the trial. DNA extraction and genotyping was co-funded by the National Institute for Health Research (NIHR), the NIHR BioResource (http://bioresource.nihr.ac.uk) and the NIHR Cambridge Biomedical Research Centre (BRC-1215-20014). Olink® Proteomics assays were funded by Biogen, Inc. (Cambridge, MA, US). SomaLogic assays were funded by Merck and the NIHR Cambridge BRC (BRC-1215-20014). The academic coordinating centre for INTERVAL was supported by core funding from: NIHR Blood and Transplant Research Unit in Donor Health and Genomics (NIHR BTRU-2014-10024), UK Medical Research Council (MR/L003120/1), British Heart Foundation (SP/09/002; RG/13/13/30194; RG/18/13/33946) and the NIHR Cambridge BRC (BRC-1215-20014). A complete list of the investigators and contributors to the INTERVAL trial is provided in reference [21]. The academic coordinating centre would like to thank blood donor centre staff and blood donors for participating in the INTERVAL trial.

This work was supported by Health Data Research UK, which is funded by the UK Medical Research Council, Engineering and Physical Sciences Research Council, Economic and Social Research Council, Department of Health and Social Care (England), Chief Scientist Office of the Scottish Government Health and Social Care Directorates, Health and Social Care Research and Development Division (Welsh Government), Public Health Agency (Northern Ireland), British Heart Foundation and Wellcome. This study was also supported by the Victorian Government’s Operational Infrastructure Support (OIS) program.

This work uses data provided by patients and collected by the NHS and Public Health England (PHE) as part of their care and support. Data on Hospital Episode Statistics, mortality and cancer registration was obtained from NHS Digital (data sharing agreement reference: DARS-NIC-156334-711SX).

S.C.R and J.M. were funded by the NIHR Cambridge BRC (BRC-1215-20014). S.A.L. is supported by a Canadian Institutes of Health Research postdoctoral fellowship (MFE-171279). G.A. was supported by a National Health and Medical Research Council of Australia (NHMRC) Early Career Fellowship (no. 1090462). A.V.K. was supported by grants from the National Human Genome Research Institute (award numbers 1K08HG010155 and 5UM1HG008895), an institutional grant from the Broad Institute of MIT and Harvard (variant2function), and a Hassenfeld Scholar Award from Massachusetts General Hospital. J.D. holds a British Heart Foundation Personal Professorship and an NIHR Senior Investigator Award.

The funders had no role in study design, data collection and analysis, decision to publish, or preparation of the manuscript. The views expressed in this manuscript are those of the author(s) and not necessarily those of the NIHR or the Department of Health and Social Care.

## Data Availability

With the exception of electronic hospital records, all data used in this study is publicly available or deposited in a public repository. Genetic data, proteomic data, and basic cohort characteristics for the INTERVAL cohort are available via the European Genotype-phenome Archive (EGA) with study accession EGAS00001002555 (https://www.ebi.ac.uk/ega/studies/EGAS00001002555). Dataset access is subject to approval by a Data Access Committee: these data are not publicly available as they contain potentially identifying and sensitive patient information. Linked electronic hospital records are currently only available to researchers at the University of Cambridge UK, however, may become more widely available in the future. Contact the data access committee for further details. All other data used in this study is publicly available without restriction. The PGS used in this study are available to download through the Polygenic Score Catalog (https://www.pgscatalog.org/) with accession numbers PGS000727 (atrial fibrillation), PGS000018 (coronary artery disease), PGS000728 (chronic kidney disease), PGS000039 (ischaemic stroke), and PGS000729 (type 2 diabetes). GWAS summary statistics used to generate new PGS in this study are available to download through the GWAS Catalog (https://www.ebi.ac.uk/gwas/) with study accessions GCST008065 (chronic kidney disease), GCST007517 (type 2 diabetes), and GCST006414 (atrial fibrillation). Summary statistics for all statistical tests are available in **Supplementary Data 3**. Full pQTL summary statistics published by Sun *et al*. 2018 for all SomaLogic SOMAscan aptamers are available to download from https://www.phpc.cam.ac.uk/ceu/proteins/. A listing of *cis*-pQTLs mapped for this study are provided in **Supplementary Data 4**. GWAS summary statistics used for Mendelian randomisation are available to download through the GWAS Catalog (https://www.ebi.ac.uk/gwas/) with study accessions GCGCST004787 (coronary artery disease), GCST008065 (chronic kidney disease), GCST006906 (ischaemic stroke) and GCST007518 (type 2 diabetes). The DrugBank database is publicly available to download at https://www.drugbank.ca/releases/latest.

## Code Availability

Code used to generate the results of this study, along with a detailed list of software and versions, are available on GitHub at https://github.com/sritchie73/cardiometabolic_PGS_plasma_proteome/ which is permanently archived by Zenodo^39^ at doi: 10.5281/zenodo.4551565.

## Online Methods

### INTERVAL cohort

INTERVAL is a cohort of approximately 50,000 participants nested within a randomised trial studying the safety of varying frequency of blood donation^20,21^. Participants were blood donors aged 18 years and older (median 44 years of age; 49% women) recruited between June 2012 and June 2014 from 25 centres across England. The collection of their blood samples for research purposes was done using standard protocols and has been extensively described previously^20^. Participants gave informed consent and this study was approved by the National Research Ethics Service (11/EE/0538).

Electronic health records were obtained for all INTERVAL participants from the National Health Service (NHS) hospital episode statistics database (https://digital.nhs.uk/data-and-information/data-tools-and-services/data-services/hospital-episode-statistics) for all events up to the 8^th^ of February 2020, prior to the onset of the COVID19 pandemic in England. The median and maximum follow-up time were 6.9 years and 7.7 years respectively. The earliest available hospital record for any INTERVAL participant was the 25^th^ March 1999, with maximum retrospective follow-up of 13.6 years. These records came in the form of international classification of diseases 10^th^ revision (ICD-10) codes^40^ and were subsequently made available to analysts after summarisation into 301 endpoints using CALIBER rule-based phenotyping algorithms^41^ (https://www.caliberresearch.org/portal). ICD-10 codes contributed to each event regardless of whether they coded for primary or non-primary diagnoses in the hospital records.

Genotyping, quality control, and imputation of INTERVAL participants has been described in detail previously^42^. Briefly, participants were genotyped using the Affymetrix UK Biobank Axiom array in 10 batches. Samples were removed if they had sex mismatch, extreme heterozygosity, were of non-European descent, or were duplicate samples. Related samples were removed by excluding one sample from each pair of close relatives (first or second degree; identity-by-descent 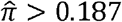). Genotyped variants were removed if they were monomorphic, bi-allelic and had Hardy-Weinberg equilibirum p-value < 5×10^−6^, or call rate < 99%. SHAPEIT3 was used to phase variants, then imputation to the UK10K/1000 Genomes panel was performed using the Sanger Imputation Server (https://imputation.sanger.ac.uk).

Quantification, processing, and quality control of protein levels in INTERVAL using the SOMAscan assays has been described in detail previously^22^. Briefly, relative concentrations of 4,034 SOMAscan aptamers were measured in 3,562 INTERVAL participants in two batches by SomaLogic Inc. (Boulder Colorado, US) using version 3 of the SOMAscan platform. Aptamers were excluded if, in the latest version of the SOMAscan platform, they (1) targeted non-human proteins, (2) have been found to be measuring the fusion construct rather than the target protein, or (3) found to be measuring a contaminant. A curated information sheet for all 4,034 aptamers is provided in **Supplementary Data 1**.

Aptamer concentrations (relative fluorescence units) were natural log transformed then adjusted within each batch for participant age, sex, the first three genetic PCs, and duration between blood draw and sample processing (< 1 day or > 1 day), then the residuals were inverse rank normal transformed. Here, we further adjusted the normalized protein levels used in previous studies for batch number, and filtered to 3,793 high quality aptamers targeting 3,438 proteins after obtaining the latest information about aptamer sensitivity and specificity from SomaLogic. Distributions of aptamer levels and associations with covariates before and after quality control are given in **Supplementary Data 2**.

In total, there were 3,087 INTERVAL participants passing quality control, without prevalent cardiometabolic disease (see below), and with matched genotype, proteomic, and electronic health record data available for the primary analyses.

### Prevalent disease exclusion

National Health Service (NHS) Blood and Transplant blood donation eligibility criteria (https://www.blood.co.uk/who-can-give-blood/) meant there were built in exclusions for the INTERVAL cohort for people with a history of major diseases, recent illness, or infection. Specifically for cardiometabolic diseases, blood donation eligibility criteria excluded individuals who had been diagnosed with atrial fibrillation, had a history of any stroke, or a history of major heart disease; including heart failure, coronary thrombosis, myocardial infarction, cardiomyopathy, ischaemic heart disease, and arrhythmia, or surgery for a non-congenital heart conditions. Use of aspirin or other blood thinners to control elevated blood pressure (hypertension) also made people ineligible to donate blood and participate in the INTERVAL cohort. Individuals with type 2 diabetes were ineligible, unless their type 2 diabetes was well controlled by diet alone, did not require regular insulin treatment, and the individual had not required insulin treatment for at least four weeks prior to attempted blood donation. Extended details on blood donation criteria eligibility for specific diseases, medications, and lifestyle factors can be found at https://my.blood.co.uk/knowledgebase.

In addition to intrinsic exclusion due to blood donation eligibility criteria, participants were excluded from analyses if they had any events relating to cardiometabolic disease prior to baseline assessment. Among the 301 CALIBER endpoints, we classified 48 as cardiometabolic disease or having potential to introduce reverse causality by modifying risk for incident AF, CAD, CKD, IS, or T2D (**Supplementary Table 1**). In total 87 participants (2.7%) were excluded, predominantly due to prevalent hypertension (N=57 events; 66% of excluded participants) and prevalent diabetes (N=11 events; 13% of excluded participants); with all others accounting for less than 5% of excluded participants (**Supplementary Table 1**).

### Polygenic scores

PGSs were derived in a consistent manner, by linkage-disequilibrium thinning, at an r^2^ threshold of 0.9, the latest GWAS summary statistics for each respective disease (**Supplementary Information**). GWAS summary statistics used to derive the AF PGS, CKD PGS, and T2D PGS were those published by Nielsen *et al*. in 2018^9^ (GCST006414), Wuttke *et al*. in 2019^10^ (GCST008065), and Mahajan *et al*. in 2018^11^ (GCST007517), respectively. PGSs for CAD and IS used in this study were our previously published CAD metaGRS^43^ and Stroke metaGRS^44^. The CAD PGS was derived from meta-analysis of three PGSs for CAD, including a PGS derived as described above from GWAS summary statistics published by Nikpay *et al*. in 2015^45^. The IS PGS was derived from meta-analysis of PGS for ischaemic stroke and its risk factors, including a PGS derived as described above from GWAS summary statistics for IS published by Malik *et al*. in 2018^12^. The PGSs each comprised 1.75–3.23 million SNPs genome-wide and are available to download through the Polygenic Score Catalog^46^ (https://www.pgscatalog.org/) with accession numbers PGS000727 (atrial fibrillation), PGS000018 (coronary artery disease), PGS000728 (chronic kidney disease), PGS000039 (ischaemic stroke), and PGS000729 (type 2 diabetes). All PGSs were derived from GWAS summary statistics including only individuals with European ancestry. See **Supporting Information** and **Extended Data Fig. 4** for details on PGS validation.

Levels of each PGS (sum of dosages × weights) were computed in INTERVAL from probabilistic dosage data using plink (version 2)^47^ after mapping PGS variants to those available in the INTERVAL genotype data (**Supplementary Information**). Levels of each PGS were adjusted for the first 10 principal components (PCs) of the imputed genotype data and standardised to have mean of 0 and standard deviation of 1 prior to downstream statistical analyses.

### PGS to protein associations

Each of the five PGSs were tested for association with each of the 3,793 aptamers using linear regression (**Fig 1a,b, Extended Data Table 2**). PGS and proteins were adjusted for covariates and normalised prior to model fitting (see above). Linear regression coefficients were averaged where multiple high quality aptamers targeted the same protein (**Supplementary Information**). False discovery rate (FDR) correction was subsequently applied across the 3,438 P-values (one per protein) for each PGS separately. Details on aptamer specificity and sensitivity are given in **Supplementary Table 2** for the 54 aptamers targeting the 49 PGS-associated proteins, and aptamer specific estimates of PGS on protein levels are detailed in **Supplementary Table 3** for the five PGS-associated proteins targeted by more than one aptamer (WFIKKN2, GPD1, IGFBP1, IGFBP2, and SHBG).

### Polygenicity of PGS to protein associations

To quantify the polygenicity of PGS to protein associations (**Fig. 1c, Extended Data Fig. 3**) we performed a multi-step experiment to determine the proportion of the genome required to explain that association. First, we split the given PGS into separate scores for each of the 1,703 approximately independent LD blocks estimated in Europeans from the 1000 Genomes reference panel by Berisa & Pickrell 2016^48^ (https://bitbucket.org/nygcresearch/ldetect-data/src/master/EUR/fourier_ls-all.bed). Next, we tested each of these 1,703 scores for association with the given protein (**Supplementary Data 3e**). Then, we retested the PGS to protein association, progressively removing independent LD blocks, at each step removing the LD block whose score had the strongest association with the protein. From this we quantified the polygenicity (**Fig. 1c**) based on the LD blocks needed to be removed from the given PGS in order to attenuate the PGS to protein association (so that the association P-value became > 0.05, **Supplementary Data 3f**) as the sum of removed LD block sizes / sum of all LD block sizes (*i*.*e*. proportion of genome removed). **Extended Data Fig. 3** shows the independent LD blocks contributing to the polygenicity of each PGS to protein association.

### Independent contributions of PGS and pQTLs to protein levels

Multivariable linear regression models were fit for each protein on PGS levels and pQTL dosages to estimate their independent contributions to protein levels (**Supplementary Table 4**). The pQTLs used for each protein were: (1) conditionally independent pQTLs mapped in INTERVAL and published by Sun *et al*. 2018^22^, which included both *cis* (within 1Mb of the encoding gene) and *trans* pQTLs passing the *trans*-significance threshold of P < 1.5×10^−11^; (2) *trans*-pQTLs with P < 1.5×10^−11^ (lead variant only) for proteins not published in Sun *et al*. 2018^22^ (B2M, DUSP26, and FTMT); and (3) hierarchically significant *cis*-pQTLs (lead variant only) mapped in this study (**Supplementary Data 4, Supplementary Information**) for proteins without *cis*-pQTLs passing the trans-pQTL significance threshold above (ACY1, ADIPOQ, APOE, CST3, GPD1, PTPRU, SHBG, and UST).

### Incident disease associations

PGSs and protein levels were tested for association with incident disease using Cox proportional hazards models adjusting for age and sex (**Fig. 2b, Extended Data Fig. 4**) using the survival package in R. The timescale used was time from baseline to first event of the relevant disease or to the latest available date in the hospital records (8^th^ February 2020). PGSs and proteins were adjusted for covariates and normalised prior to model fitting (see above). Cox model coefficients were averaged where multiple high quality aptamers targeted the same protein (**Supplementary Information**).

Incident disease events for AF, CAD, CKD, IS, and T2D were defined as first hospital episode for the closest matching CALIBER phenotype^41^ (https://www.caliberresearch.org/portal). Incident AF events were defined as any hospital episode with ICD-10 code I48. Incident IS events were defined as any hospital episode with ICD-10 codes I63 or I69.3. For CAD we analysed incident MI events, defined as any hospital episode with ICD-10 codes I21–I23, I24.1, or I25.2. The closest matching CALIBER phenotype for T2D was for diabetes more broadly, including ICD-10 codes for any hospital episode for type 1 or type 2 diabetes or complications thereof: E10–E14, G59.0, G63.2, H28.0, H36.0, M14.2, N08.3, or O24.0–O24.3, however we note type 1 diabetics are not eligible to donate blood (https://my.blood.co.uk/knowledgebase/) and adult onset of type 1 diabetes is relatively rare compared to type 2 diabetes^49^. closest matching CALIBER phenotype for CKD was for end stage renal disease more broadly, which as defined as any hospital episode with ICD-10 codes N16.5, N18.5, T82.4, T86.1, Y60.2, Y61.2, Y84.1, Z49.1, Z49.2, Z94.0, and Z99.2.

### Mediation analysis

Mediation analysis was used to identify causal proteins by identifying the PGS-associated proteins which partially mediate the association of PGS on disease (**Fig. 2d**). This approach uses the counterfactual framework to infer causal effects ^28,50,51^ and can be adapted to this setting as the arrow of causality between PGS and any associated phenotype can only flow in one direction as the PGS is fixed at conception (i.e. the underlying alleles in each person cannot be modified later in life by protein levels or the development of cardiometabolic disease). Here, we used the natural effects model developed by Vansteelandt *et al*. 2012^52^, which is available in the medflex R package^53^, to estimate natural indirect effects (effects of PGS on disease through protein levels) on the log odds scale by imputing unobserved counterfactuals. Standard errors were computed using the robust sandwich estimator^54^, from which 95% confidence intervals and P-values were calculated. Multiple mediation analysis^55^ was performed using the R package mma^56^ to quantify the proportion of PGS to disease association mediated by the 15 causal T2D proteins.

### Mendelian randomisation

Two-sample Mendelian randomisation^29^ was also performed as an orthogonal approach to identify proteins which may play a causal role in disease (**Extended Data Fig. 5, Supplementary Table 5**,**6**). PGS-associated proteins were tested provided they had three or more independent by LD (r^2^ < 0.1) *cis*-pQTLs after mapping pQTL to GWAS summary statistics (**Supplementary Information**), and provided the SomaLogic aptamer(s) did not have similar affinity for or comparable binding to multiple proteins or differential binding to specific isoforms (**Supporting Table 3, Supplementary Information**). In total, 12 of the 49 PGS-associated proteins could be tested (24%), substantially higher than the overall measured proteome (497 proteins, 14.5%). GWAS summary statistics were obtained from Nelson *et al*. 2017^13^ for coronary artery disease (GCST004787), Wuttke *et al*. 2019^10^ for chronic kidney disease (GCST008065), Malik *et al*. 2018^12^ for ischaemic stroke (GCST006906) and Mahajan *et al*. 2018^11^ for type 2 diabetes (GCST007518). In all cases, we used the GWAS summary statistics for the samples of recent European ancestry. For type 2 diabetes, we used the BMI-adjusted GWAS summary statistics in order to avoid false positive causal estimates arising where pQTLs influence type 2 diabetes risk through BMI rather than through the tested protein (horizontal pleiotropy). We used five different Mendelian Randomisation methods^57–60^, each of which make use of information across 3 or more instruments to estimate causal effects with each method differentially robust to different sources of bias, to obtain a consensus (median) estimate of causal effects of protein levels on disease risk (**Supporting Information**). We considered there to be a significant causal effect where P < 0.05 along with no significant evidence that causal effects were due to associations of the pQTLs with some other causal risk factor (horizontal pleiotropy; Egger intercept^60^ P > 0.05). FDR correction was performed across all tested proteins for each disease separately. Analysis was performed using the R package MendelianRandomization^61^. Colocalisation analysis^62^ was also performed where *cis*-pQTL instruments had P < 1×10^−6^ in the respective GWAS (**Supplementary Table 6, Supplementary Information**).

### Drug targets

For each PGS-associated protein, a list of drugs that target or interact with the protein was downloaded from the DrugBank database^30^ version 5.17 released on the 2^nd^ of July 2020 (https://go.drugbank.com/releases/latest) (**Extended Data Table 5, Supplementary Table 7**). To obtain a list of drugs that counteract PGS effects and thus may have potential repurposing opportunities (**Table 1**), we filtered to drugs with approved status and not withdrawn status, then to drugs whose effect on the protein was in the opposite direction to the effect of the PGS on protein levels (e.g. inhibitors where increased PGS was associated with increased protein levels, **Supplementary Information**).

## Extended Data

**Extended Data Figure 1:**
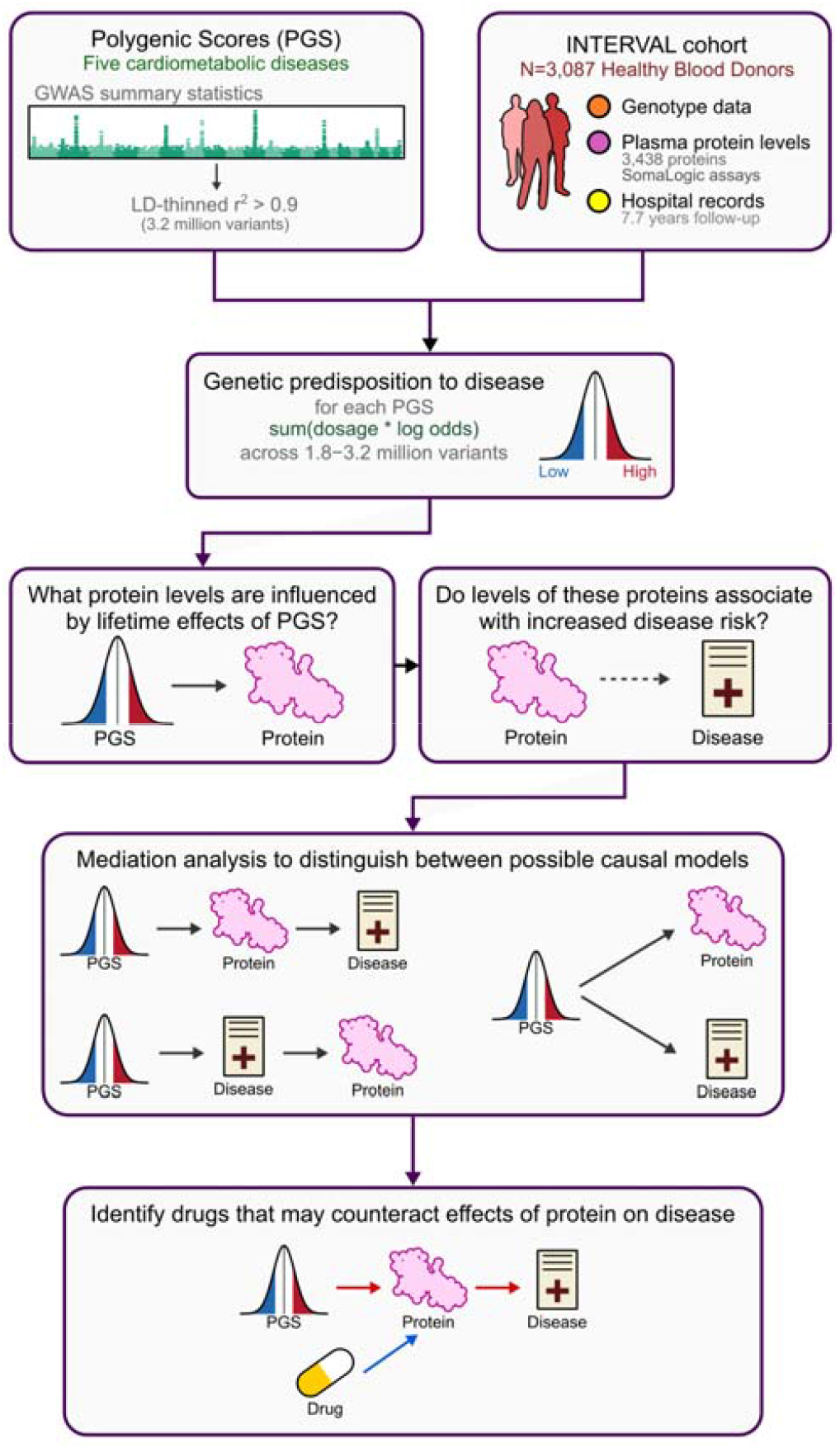
Study schematic.

**Extended Data Table 1:**
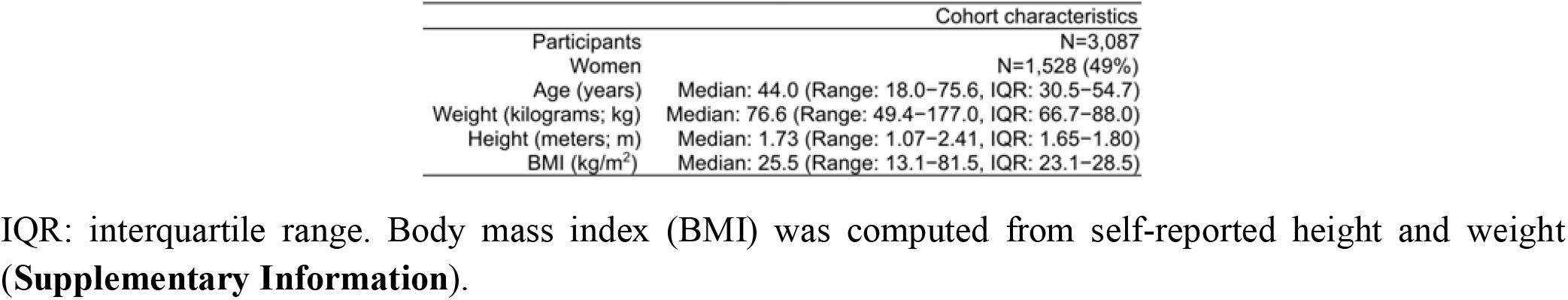
Cohort characteristics.

**Extended Data Table 2:**
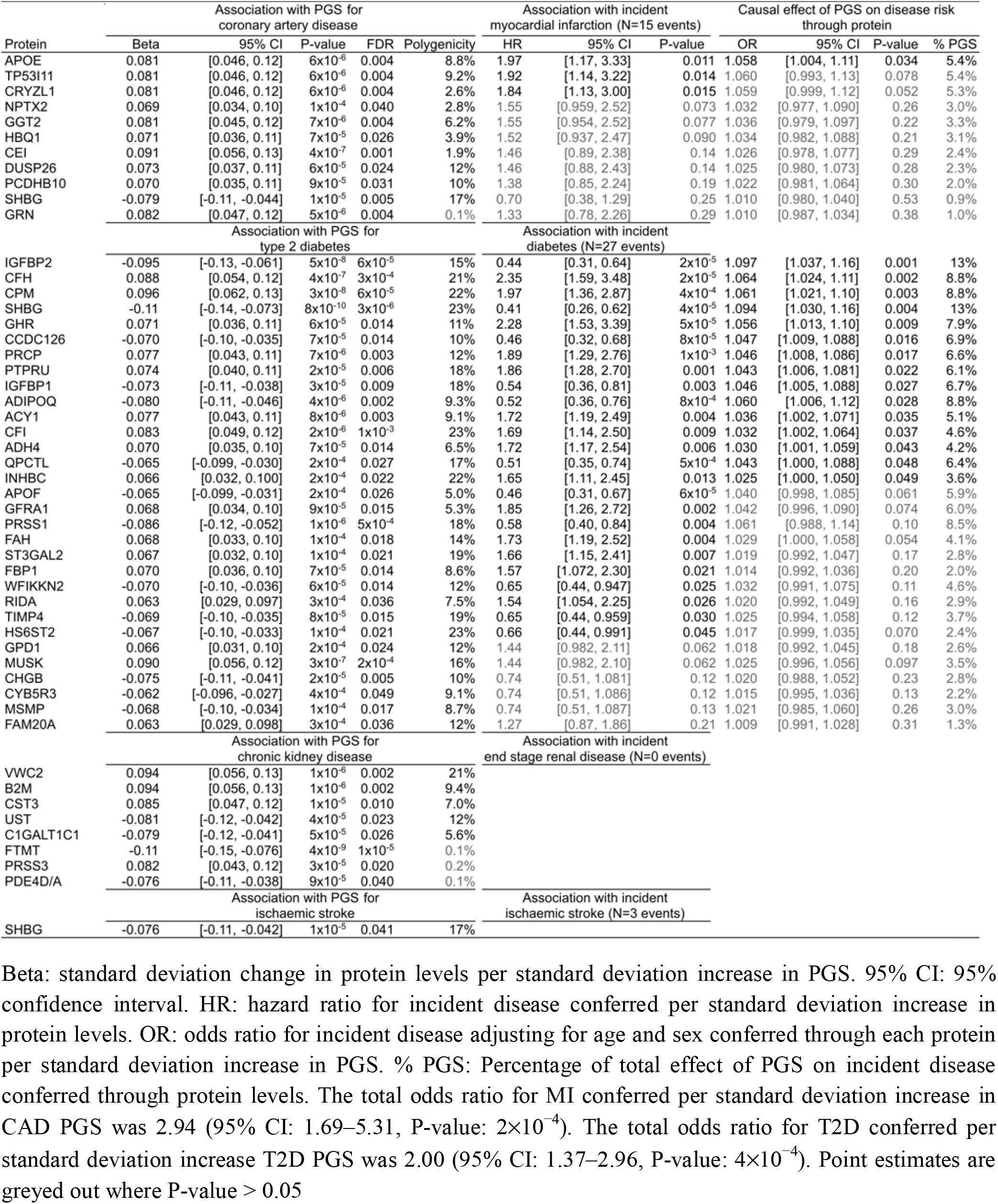
Point estimates for PGS to protein to disease associations.

**Extended Data Table 3:**
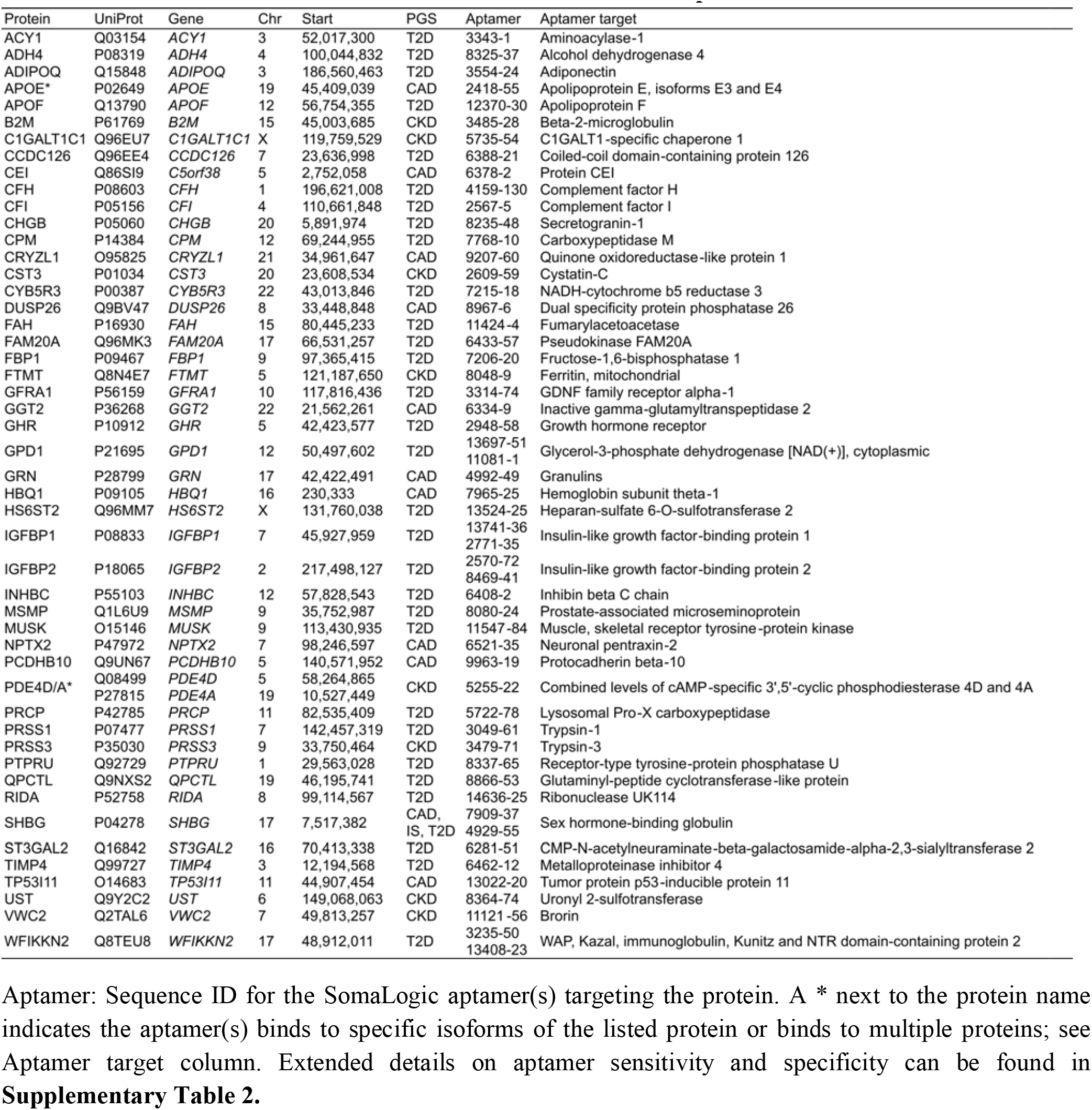
Information about each PGS associated protein.

**Extended Data Table 4:**
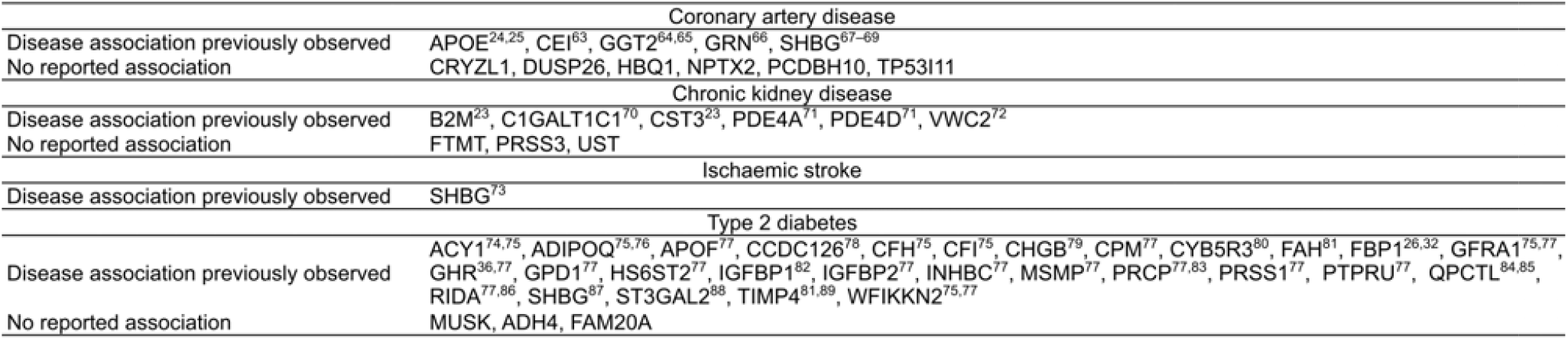
Previous evidence for PGS-associated proteins in disease.

**Extended Data Figure 2:**
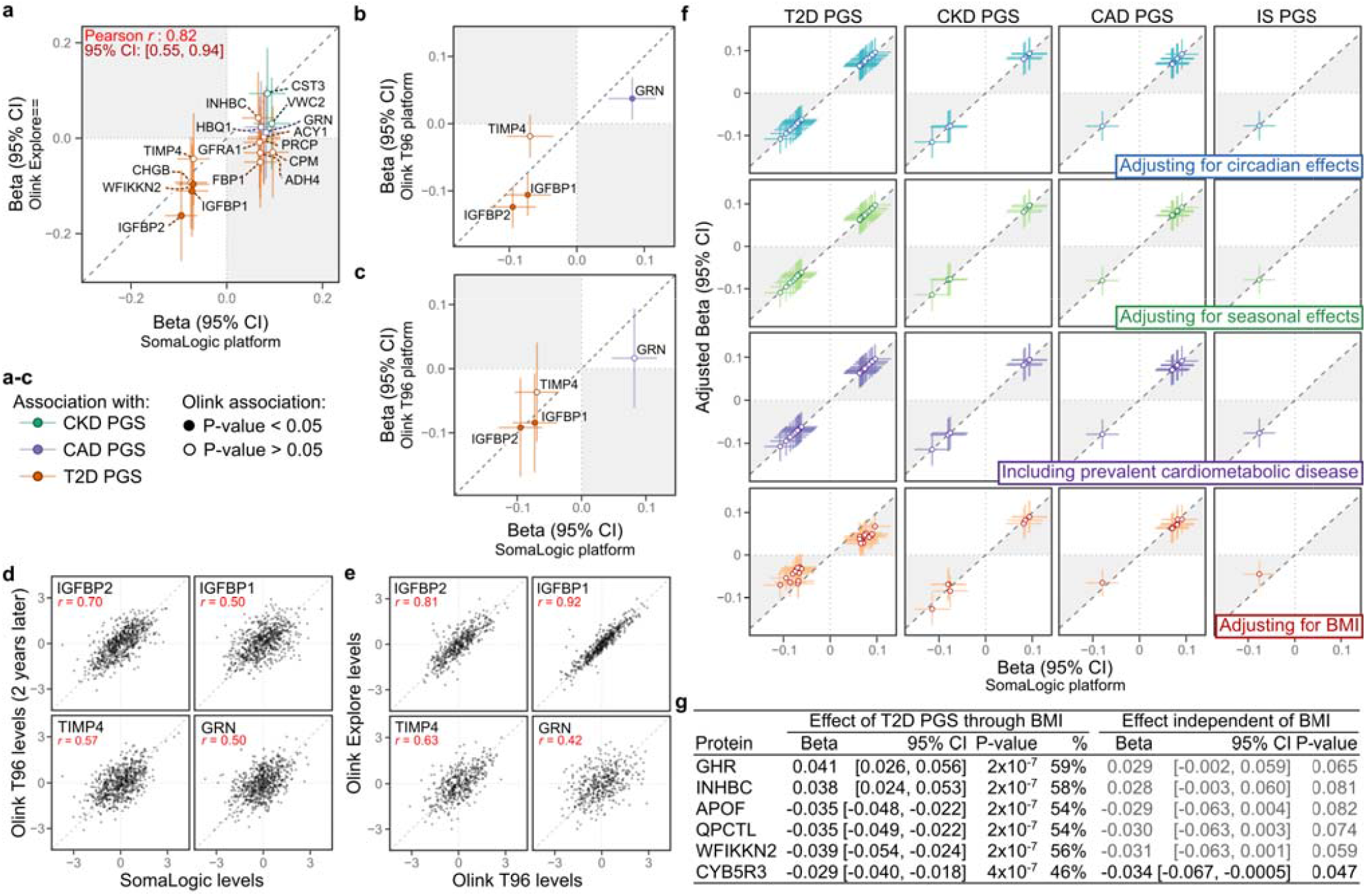
Robustness of PGS to protein associations. **a-c)** Robustness of PGS to protein associations to proteomics technology. **c)** Longitudinal stability of PGS to protein associations. **d)** Longitudinal stability of protein levels. **d**-**e)** Robustness of protein levels to proteomics technology. **f)** Robustness of PGS to protein associations to environmental and physiological confounding. **g)** Mediation of PGS to protein associations through body mass index (BMI) for six proteins associated with PGS for type 2 diabetes. **a)** Compares associations between PGSs and protein levels quantified by SomaLogic SOMAscan aptamers (x-axis; **Fig. 1b**) to associations with protein levels quantified using the Olink Explore platform in 418 independent INTERVAL participants (y-axis) with no prevalent cardiometabolic disease (**Supplementary Information**). In total 1,463 proteins were quantified by the Olink Explore platform, including 907 quantified by the SomaLogic platform, and among these 16 of the 49 PGS-associated proteins. Points correspond to PGS to protein level association beta estimates, and the bars to their 95% confidence intervals. **b)** Compares associations between PGSs and protein levels quantified by SomaLogic SOMAscan aptamers (x-axis; **Fig. 1b**) to associations with protein levels quantified using the Olink T96 platform in 3,848 independent INTERVAL participants. In total 265 proteins were quantified by the Olink T96 platform, including 224 quantified by the SomaLogic platform, and among these, 4 of the 49 PGS-associated proteins. **c)** Compares PGS to protein associations in 646 participants with protein levels quantified by both the SomaLogic platform and Olink T96 platform (from blood samples taken after 2 years of follow-up). **a-c**) share common x-axes and legend. Point estimates for associations between PGS and protein levels assessed by Olink proteomics in each panel are given in **Supplementary Data 3b. d)** Compares protein levels quantified by the SomaLogic platform (x-axes) to protein levels quantified by the Olink T96 platform (y-axes) after two years of follow-up in 646 participants. **e)** Compares protein levels quantified by the Olink T96 platform (x-axes) to protein levels quantified by the Olink Explore platform (y-axes). **f)** Compares PGS to protein associations before (x-axes; **Fig. 1a**) and after (y-axes) adjustment for circadian effects (time of day of blood draw), adjustment for seasonal effects (date of blood draw), when including 87 participants with prevalent cardiometabolic disease, and adjustment for BMI. To capture the potentially non-linear effects of circadian rhythm and season on protein levels both were treated as categorical variables with 10 groups of equal length duration, using the group with the largest sample size as the reference in the model (**Supplementary Information**). Point estimates in sensitivity analyses are given in **Supplementary Data 3c. g)** For the six proteins whose association T2D PGS was attenuated (P > 0.05; **Extended Data Fig. 2f**) gives, from mediation analysis (**Online Methods**), the estimated effect of T2D PGS on the protein levels through BMI (standard deviation change in protein levels through BMI per standard deviation increase in T2D PGS) and the estimated effect of T2D PGS on protein levels independent of BMI. 95% CI: 95% confidence interval.

**Extended Data Figure 3:**
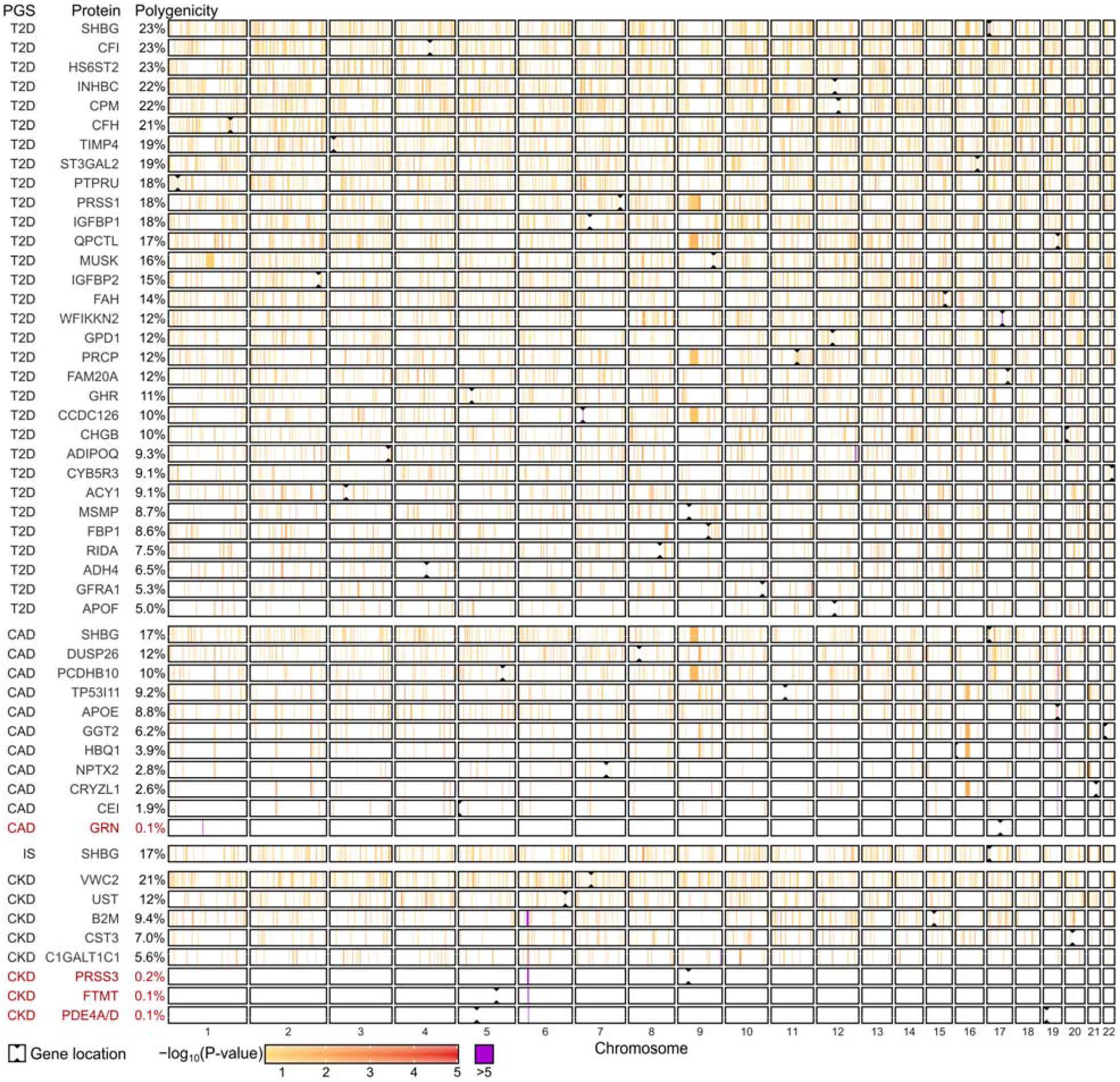
Polygenicity of PGS to protein associations. Linkage disequilibrium (LD) blocks contributing to each PGS to protein association (**Online Methods**). Each PGS was partitioned into 1,703 approximately independent LD blocks^48^ then tested for association with each protein (**Supplementary Data 3e**). To obtain the set of LD blocks contributing to each PGS to protein association, LD blocks were removed from the PGS in ascending order by association P-value until the PGS to protein association was attenuated (P > 0.05; **Supplementary Data 3f**). Here, associations (-log_10_ P-values) between protein levels and LD blocks contributing to the PGS to protein association are shown. Regions in white contain LD blocks that did not contribute to the PGS to protein association. The total percentage of the genome contributing to the PGS to protein association (polygenicity; **Fig. 1c**) is shown on the right. PGS to protein associations listed in red are those explained by pQTLs (*cis* and/or *trans*) rather than polygenic.

**Extended Data Figure 4:**
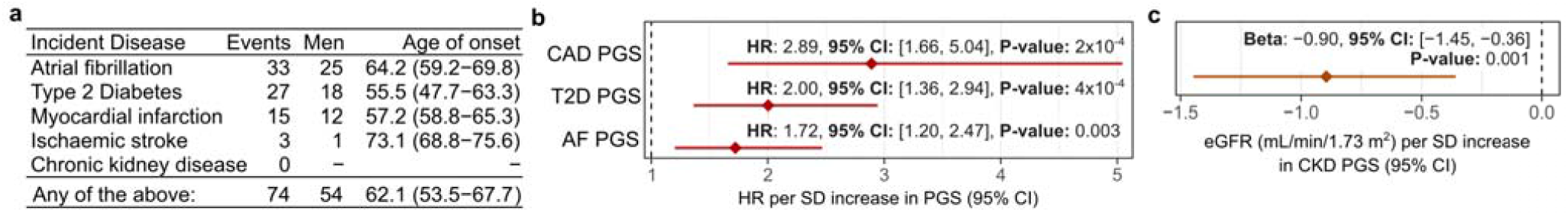
Incident disease and PGS validity. **a)** Incident disease events over the 7.7 year of follow-up in the 3,087 INTERVAL participants. Endpoint: incident disease definition available in INTERVAL for the relevant PGS, as defined by CALI ER phenotyping algorithms (**Online Methods**). Age of onset: median age of first hospitalisation with the respective endpoint. Numbers in brackets gives the interquartile range. **b)** Hazard ratio (HR) and 95% confidence interval (95% CI) conferred per standard deviation increase of the respective PGS on risk of hospitalisation with the relevant endpoint. CAD PGS was tested for incident myocardial infarction. Hazard ratios were fit using cox proportional hazards models, adjusting for age and sex, and 10 genetic PCs. **c)** Association between PGS for chronic kidney disease with estimated glomerular filtration rate (eGFR), a marker of renal function used in chronic kidney disease diagnosis (**Online Methods**): decreased eGFR is indicative of reduced renal function. EGFR was computed from serum creatinine in 3,307 participants using the CKD-EPI equation (**Supplementary Information**). Association was fit with linear regression adjusting for age and sex, and 10 genetic PCs.

**Extended Data Figure 5:**
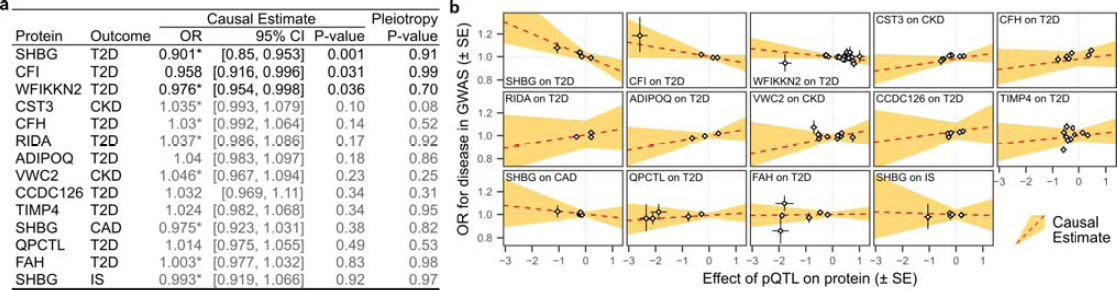
Mendelian randomisation analysis. **a)** Causal effects of protein levels on disease risk estimated through two-sample Mendelian randomisation analysis of pQTL summary statistics and disease GWAS summary statistics (**Online Methods**). OR: consensus estimate of the odds ratio conferred per standard deviation increase in protein levels across five Mendelian randomisation methods (**Supplementary Information**; **Supplementary Table 5**). * Estimated causal effect is directionally consistent with PGS to protein to disease associations in **Fig. 2**. 95% CI: 95% confidence interval. Pleiotropy P-value: P-value for the intercept term in Egger regression, which indicates where P < 0.05, confounding of the causal estimate by associations between genetic instruments (*cis-*pQTLs) with multiple disease risk factors (horizontal pleiotropy). Entries are greyed out where P > 0.05. **b)** Dose response curves showing the estimated causal effect of changes in protein levels on disease risk for each protein and disease. The slope of the orange dashed line corresponds to the estimated causal effect (Odds Ratio from **Extended Data Fig. 5a**). The yellow ribbon shows the 95% confidence interval for the estimated causal effect (slope), accounting also for the 95% confidence interval for the intercept term in Egger regression. Points on each plot show the *cis*-pQTLs used as genetic instruments for each test (**Supplementary Table 6**). On the x-axes, points show the standard deviation change in protein levels per copy of the minor allele, and horizontal bars indicate the standard error. On the y-axes, points show odds ratio conferred per copy of the minor allele, and vertical bars indicate the standard error.

**Extended Data Table 5:**
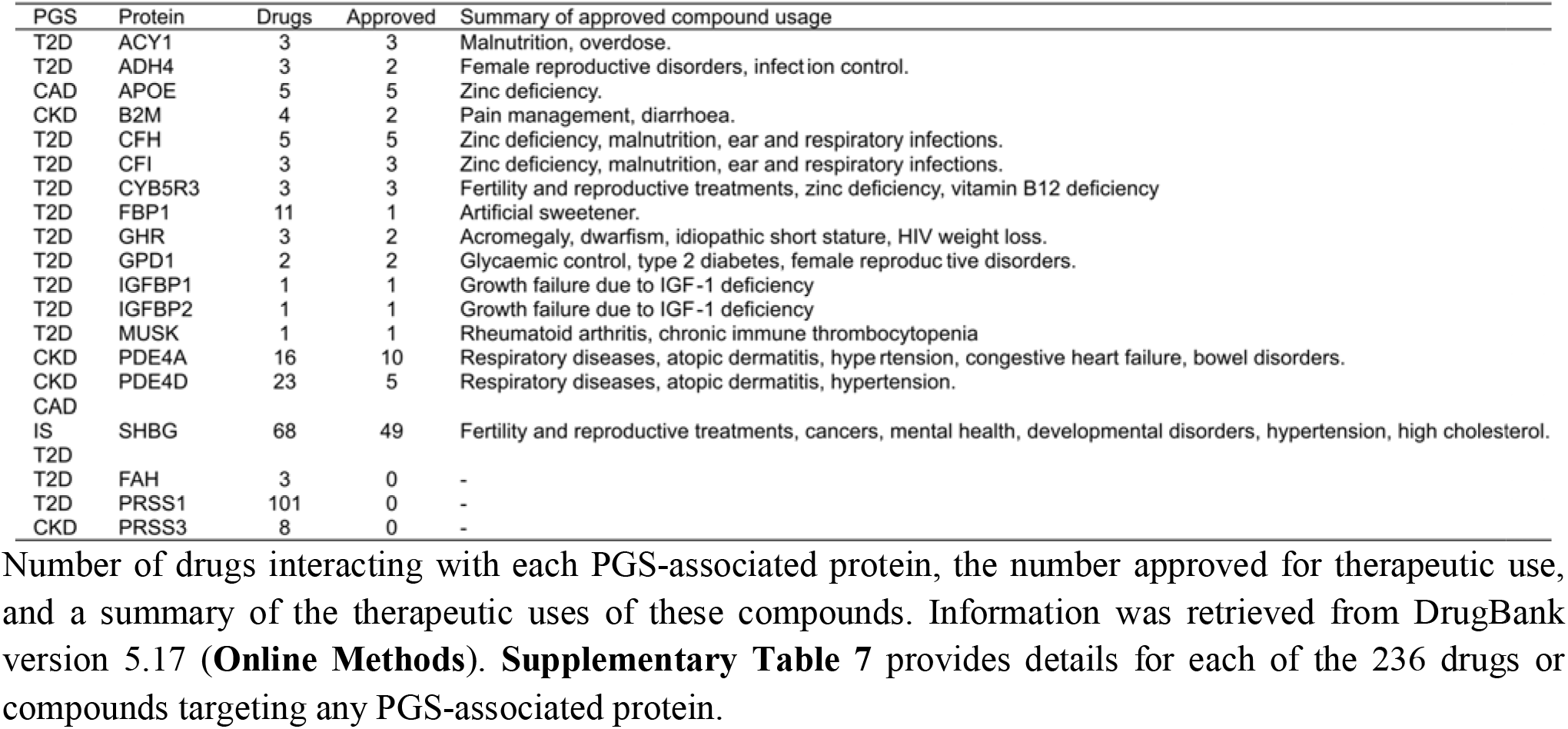
PGS-associated drug targets.

