## Supplementary Information for "Integrative analysis of the plasma proteome and polygenic risk of cardiometabolic diseases"

### Supplementary Notes

#### PGS validation

The CAD PGS and IS PGS have been externally validated in the studies in which they were published^43,44^ (see also their respective pages on the PGS Catalog, which also includes external validation by other independent studies, using their accession numbers PGS000018 and PGS00039 at https://www.pgscatalog.org/). The AF PGS, CKD PGS, and T2D PGS were derived *de novo* for this study (**Online Methods**). In each case, PGS derivation did not include INTERVAL participants, and thus was able to act as an external validation cohort. Here, we show strong association between AF PGS and 7.7 year risk of AF (Hazard Ratio: 1.72, 95% Confidence Interval: 1.20–2.47, P-value: 0.003; **Extended Data Fig. 4b**) and between T2D PGS and 7.7 year risk of T2D (Hazard Ratio: 2.00, 95% Confidence Interval: 1.36–2.94, P-value: 4×10^-4^; **Extended Data Fig. 4b**) in Cox proportional hazard models adjusting for age and sex (**Online Methods**). There were insufficient events to test association between CKD PGS and 7.7 year disease risk (N=0 events; **Extended Data Fig**. **4a**), however, we observed strong correlation between the CKD PGS and eGFR (Beta: −0.90 standard deviation change in eGFR per standard deviation increase in CKD PGS, 95% confidence interval: −1.45– −0.36, P-value: 0.001; **Extended Data Fig. 4c**). Decreased eGFR indicates decreased renal function, and is the primary method through which CKD is diagnosed in the clinic^90^.

#### Pathway enrichment

No pathways, gene sets, or protein-protein interaction networks were significant after multiple testing correction (**Supplementary Methods**). An archive of the query can be found at https://biit.cs.ut.ee/gplink/l/aWke5_FsQ1 for the proteins associated with the CAD PGS, at https://biit.cs.ut.ee/gplink/l/UNpRmoI8QF for the proteins associated with the CKD PGS, and at https://biit.cs.ut.ee/gplink/l/3QIUMijpTa for the proteins associated with the T2D PGS.

#### Robustness to proteomics technology

To test the robustness of PGS to protein associations, we measured 1,463 proteins in 418 independent samples with no prevalent cardiometabolic disease using an orthogonal and complementary technology; the Illumina NovaSeq based Olink Explore quantitative protein assays (**Supplementary Methods**). These allowed us to independently test 16 proteins with significant PGS to protein associations, showing that effect size estimates were strongly correlated between platforms (**Extended Data Fig. 2a**; Pearson *r*: 0.82, P-value: 9×10^-5^). We also tested four of these proteins using the qPCR based Olink Target 96 (Olink T96) protein assays in 3,848 independent samples (**Supplementary Methods**), where effect size estimates were again similar and three of four PGS associations remained statistically significant (**Extended Data Fig. 2b**).

#### Longitudinal stability

In a further 646 samples with SomaLogic protein data at baseline and Olink T96 protein data in samples taken two-years after baseline, we observed similar effect size estimates for these four PGS to protein associations (**Extended Data Fig. 2c**) with strong correlations in protein levels across time and between platforms (Pearson *r*: 0.50–0.70; **Extended Data Fig. 2d**), further indicating temporal stability of the effects of PGS on protein levels.

#### Mendelian randomisation

Mendelian randomisation analysis found significant evidence (consensus causal estimate P-value < 0.05 and horizontal pleiotropy P-value > 0.05; **Online Methods**) for causal effects on type 2 diabetes risk for three proteins on SHBG, CFI, and WFIKKN2 (**Extended Data Fig. 5**). Causal effects for SHBG and CFI, but not for WFIKKN2, were also identified by mediation analysis (**Fig. 2d**). Directions of causal effect were consistent with associations with T2D PGS and incident T2D for SHBG and WFIKKN2 (**Extended Data Table 2**), and consistent with causal effects estimated by previous studies^75,77,87^. The direction of causal effect for CFI estimated by Mendelian randomisation was discordant with the direction of association between CFI, T2D PGS, and incident T2D (**Extended Data Table 2**), however, such discrepancies between causal effects estimated through Mendelian randomisation and direction of association in observational studies has been previously observed^22,91^ and has been postulated to arise due to time-varying effects of genetic variants on protein levels^92,93^.

#### Evidence supporting drugs for countering lifetime risk conferred by PGS

Metformin (DB00331) is a drug used to manage T2D^31,94^, which reduces liver glucose production by inhibition of FBP1^32^. Consistent with this, here we observed an association between increased FBP1 levels and increased PGS for type 2 diabetes (**Fig. 1b**) as well as increased risk of 7.7 year risk of incident T2D (**Fig. 2b**). Additionally, metformin is also known to inhibit GPD1^95^ (**Table 1**), which likewise was associated with increased T2D PGS (**Fig. 1b**).

FBP1 is also inhibited by adenosine phosphate (DB00131), which acts as an antagonist and inhibitory allosteric modulator on FBP1^96,97^. Also known as adenosine monophosphate (AMP), it is a nucleotide component of RNA that plays essential roles in cellular metabolism and intercellular signalling^98^.

Iloprost (DB01088) is a drug used to treat pulmonary arterial hypertension^99^. It is known to induce expression cAMP phosphodiesterase genes, including *PDE4D* and *PDE4A*^100–102^. Here, combined protein levels for PDE4D and PDE4A were decreased in individuals with elevated CKD PGS (**Fig. 1b**), suggesting a potential repurposing opportunity for iloprost for CKD prevention or treatment (**Table 1**). Supporting this, in phase II clinical trial (NCT00345501) iloprost has been found to prevent contrast-induced nephropathy^103^, a complication of intravenous contrast administration defined by impairment of kidney function^104,105^. Further, individuals with recessive PDE4D deletion have been found to be at increased risk for CKD^71^.

Fostamatinib (DB12010), also known as syk inhibitor R406^106^, is a drug recently approved for treatment of chronic immune thrombocytopenic purpura^107^. It is known to inhibit muscle skeletal receptor tyrosine-protein kinase (MUSK)^106^. Here, increased MUSK levels were associated with increased T2D PGS (**Fig. 1b**), suggesting a potential repurposing opportunity for fostamatinib for T2D prevention or treatment (**Table 1**).

Zinc chloride (DB14533) and zinc sulfate (DB14548) are used as nutritional supplements in intravenous nutrition to prevent complications associated with zinc deficiency^108,109^. Zinc has been found to affect the proteolysis of Apolipoprotein E (ApoE) in an isoform dependent manner, increasing proteolysis for ApoE isoform ε4^110^. Here, increased ApoE levels (isoforms ε3 and ε4) were associated with increased CAD PGS (**Fig. 1b**), increased 7.7 year risk for incident MI (**Fig. 2b**), and that these changes likely play a causal role in increasing MI risk (**Fig. 2d**). ApoE isoform ε4 is known to increase atherosclerosis risk by increasing plasma low density lipoprotein (LDL) levels^111^ and there is some evidence that elevated serum zinc levels may confer a protective effect on cardiovascular disease risk^112^. Together, these suggest dietary zinc supplementation may reduce risk of coronary artery disease.

Sex hormone binding globulin (SHBG) is a protein that binds to, carriers, and modulates the function of androgens and oestrogens and plays a role in fertility and reproduction^113–115^. Here, decreased SHBG levels were associated with increased PGSs for CAD, T2D, and IS (**Fig. 1b**), and that these changes causally led to increased 7.7 year risk for incident T2D (**Fig. 2b,d**). Further, previous studies have also shown decreased SHBG is causal for increased T2D risk^77,87^, and in women, is associated with increased risk of stroke^73^ and other cardiovascular diseases^68^. Together, these suggest that drugs which increase the levels or function of SHBG may be suitable for repurposing for treatment or prevention of T2D, and potentially also for CAD, and IS.

Three drugs that increase the levels or function of SHBG were identified: progesterone (DB00396), tamoxifen (DB00675), and ketoconazole (DB01026) (**Table 1**). Progesterone is a naturally occurring female hormone that is used in a female fertility treatments and oral contraceptives^116,117^ and is a potentiator of SHBG^118,119^. Clinical trials have investigated the use of progesterone for reducing cardiovascular disease risk factors including type 2 diabetes (NCT00000466, NCT00361075), however, evidence supporting its use is unclear due to the differences in effects for progesterone and other hormone replacement therapies between men and women, age, pre- or post- menopausal state, drug design, and delivery mechanism^120–124^ . Tamoxifen is a drug which has been used to treat oestrogen positive breast cancers^125^, and is an inducer of SHBG^119^. Clinical trials have investigated the effect of tamoxifen on cardiovascular disease risk, but have found no effect^126,127^. Ketoconazole is an oral antifungal^128^ which is carried by and acts as a ligand on SHBG^129^. It has shown efficacy in phase 2 clinical trial (NCT00494663) for treating elevated blood glucose^130^, however, has not been taken forward due to its sub-optimal safety profiles among other clinical limitations^128,130^

### Supplementary Methods

#### Polygenic score derivation

PGSs for AF, CKD, and T2D were derived by filtering their respective GWAS summary statistics (**Online Methods**) to a high-confidence set of 2.3 million common variants LD-thinned at r^2^=0.9 threshold in the UK Biobank version 3 genotype data^131,132^ (imputed to the 1000 Genomes, UK10K, and haplotype reference consortium (HRC)^133^ panels). This variant set included only bi-allelic SNPs with minor allele frequency > 1% and imputation INFO score > 0.4, and excluded SNPs which could not reliably be matched between datasets due to strand ambiguity (A/T and G/C SNPs).

#### Mapping variants between datasets

Variants in PGSs or in GWAS summary statistics (for Mendelian randomisation; see below) were mapped to variants in the INTERVAL genotype data by genomic location and effect allele (or by rsid if genomic location not provided). If a variant matched by genomic location but the effect allele matched no alleles in the genotype data, the strand orientation of the alleles were flipped and checked for match again. Variants with potential for allele mismatch due to strand ambiguity (e.g. palindromic A/T and G/C variants) were excluded unless effect allele frequency information was provided in the PGS or GWAS summary statistics. Where effect allele frequency was provided, alleles were matched additionally by frequency (yes/no below 50%), and excluded where the minor allele frequency was greater than 42%.

When mapping GWAS summary statistics to INTERVAL for Mendelian randomisation (see below), multi-allelic variants were removed prior to the procedure described above, and effect alleles were oriented to the minor allele in INTERVAL.

#### Estimated Glomerular Filtration Rate

Serum creatinine was quantified in 3,307 INTERVAL participants by Metabolon HD4 metabolomics in mg/dL units, and adjusted for sample measurement batch, sample measurement plate, and days between blood draw and sample processing. Subsequently, eGFR was quantified from serum creatinine using the CKD-EPI equation^134^:

$$eGFR=141\times{min\left( {SCr}/k, 1 \right)}^{\alpha}\times{max\left( {SCr}/k, 1 \right)}^{-1.209}\times{0.993}^{Age}\times\left[ 1.018 if Female \right]\times[1.159 if Black]$$

Where *SCr =* serum creatinine in mg/dL. For men, *k* = 0.9 and α = −0.411, and for women, *k* = 0.7 and α = −0.329. All participants analysed were white British.

#### Averaging of association summary statistics across aptamers

Many proteins were targeted by more than one SomaLogic aptamer. For these proteins, each aptamer may target different components of the protein, with varying degrees of measurement error. Throughout, multiple aptamers targeting the same protein have been treated as independent measurements, and coefficients from statistical models have been averaged to obtain a single robust estimate for the respective protein. For linear coefficients (e.g. linear regression) the mean of the beta coefficient and 95% confidence intervals was taken across the multiple aptamers. For exponent coefficients (e.g. hazards and odds ratios) a log transform was applied prior to the mean, and exponent taken of the result.

To obtain an average of P-values across aptamers, the P-values were converted to Z-scores, the mean of Z-scores computed, then converted back into a P-value. To ensure correct averaging of P-values from discordant directions of effect, two-sided P-values were converted to one-sided P-values with the alternate hypothesis chosen based on the sign of the beta coefficient (or log transformed coefficient in the case of hazard ratio or odds ratio) prior to Z-score conversion, then the resulting Z-score converted back to a two-sided P-value.

#### Pathway enrichment

We used the gProfiler tool^135^ (<https://biit.cs.ut.ee/gprofiler/gost>) to test whether PGS to protein associations were enriched for known pathways (KEGG, Reactome, WikiPathways), gene ontology (GO) terms (biological process, molecular function, and cellular compartment), or protein-protein interaction networks (CORUM, Human Protein Atlas). The set of 3,424 UniProtKB identifies covering all measured proteins and protein complexes was used as the reference (background) set for each test.

#### Olink protein quantification and quality control

Quantification of protein levels using Olink (Olink Bioscience, Uppsala, Sweden) proximity extension assays^136^ (**Extended Data Fig. 2a-e**) was performed using two complementary technologies: Olink’s Target 96 panels (Olink T96) (<https://www.olink.com/>) and Olink’s Explore 1536 platform (Olink Explore) (<https://www.olinkexplore.com/>). Participants selected for Olink measurement were selected from among those at least 50 years old at baseline, and Olink measurement was performed on plasma samples taken 2 years after baseline assessment. After excluding participants with prevalent cardiometabolic disease at this follow-up time-point, there were 4,494 participants: 646 also with SomaLogic proteomics at baseline (**Extended Data Fig. 2c,d**), and 3,848 participants without SomaLogic proteins at baseline (**Extended Data Fig. 2b**).

Quantification of proteins using the qPCR-based Olink T96 platform was performed in 5,000 INTERVAL participants using Olink’s cardiovascular II, cardiovascular III, and inflammation panels (<https://www.olink.com/>). In total 265 proteins were quantified by the three Olink T96 panels, of which 224 were also quantified on the SomaLogic platform, among which four were associated with any PGS (**Extended Data Fig. 2b-e**). All four of these proteins were quantified by the Olink T96 cardiovascular III panel. For this panel, protein levels on the log_2_ scale were regressed on age, sex, sample measurement plate, days between blood draw to sample processing, and season, then the residuals were inverse rank normal transformed.

Quantification of proteins using the Illumina NovaSeq-based Olink Explore platform was performed in 500 INTERVAL participants with Olink T96 data (**Extended Data Fig. 2a,e**). In total 1,463 proteins were quantified. Samples in the first column of each plate (N=50) were removed due to quality issues. No participants with Olink Explore data had SomaLogic protein data at baseline. Protein levels on the log_2_ scale were regressed on age, sex, sample measurement plate, and days between blood draw to sample processing, then the residuals were inverse rank normal transformed.

#### Physiological and environmental confounders

Sensitivity analysis were performed by additionally adjusting the PGS to protein aptamer linear regression models for BMI, circadian effects, and seasonal effects separately (**Extended Data Fig. 2f**).

BMI was calculated from self-reported weight and height. Fifteen samples with outlier BMI were removed by excluding samples with self-reported weight < 50 kg or > 160 kg; weights outside the range for NHS blood donation eligibility criteria (https://my.blood.co.uk/knowledgebase/Index/W), and samples with self-reported height < 1.47 m or > 2.1 m; clinical thresholds for dwarfism and gigantism respectively. After outlier exclusion the range of participant BMI was 17.0–55.8 (median 25.5). BMI was subsequently log transformed prior to model fitting. Mediation analysis was performed as described in the **Online Methods** for six PGS to protein associations attenuated by BMI (**Extended Data Fig. 2g**), using BMI as the mediator and protein levels as the outcome.

To capture the potentially non-linear effects of circadian rhythm and season on protein levels, both were treated as a categorical variables by grouping samples into 10 equal duration bins, with the largest sample size bin acting as the reference group in the model. To model circadian effects, the time of day of sample draw was split into 10 bins, each 73.5 minutes in length, with sample sizes ranging from 62 to 405 (median: 267, interquartile range: 237–325) samples. Bin six, covering 2:08pm–3:21pm with 405 samples, was used as the reference group. Samples with no time of blood draw recorded (n=468; 15%) were excluded when adjusting for circadian effects. To model seasonal effects, the date of blood draw was split into 10 bins, each 50 days in length, with sample sizes ranging from 117 to 446 (median: 283, interquartile range: 274–396) samples. Bin four, covering the 9^th^ November 2012 – 30^th^ December 2012 with 446 samples, was used as the reference group.

#### Cis-pQTL mapping

Previous pQTL mapping studies have focused on the identification of pQTLs genome-wide, requiring conservative P-value thresholds meaning many *cis*-pQTLs have been missed. Here, we applied a hierarchical multiple testing procedure^137^ that has shown to best control false discovery rates in *cis*-eQTL mapping^138^ to pQTL summary statistics for each SomaLogic aptamer published by Sun *et al.* 2018^22^. First, (1) P-values within 1Mb of any gene encoding the target protein were Bonferroni corrected for the number of tests within the respective *cis* window(s) to obtain locally corrected P-values, then (2) the smallest locally corrected P-value was taken for each of the 3,793 aptamers, then (3) FDR correction was applied across all aptamers to obtain a single globally corrected P-value for each aptamer, then (4) globally corrected P-values were filtered at FDR < 0.05 to identify aptamers with *cis*-pQTLs, finally (5) the locally corrected P-value corresponding to the global FDR = 0.05 threshold (locally corrected P = 0.0124) was used as a significance threshold to identify significant *cis*-variants for each of the aptamers with *cis*-pQTLs. *Cis*-pQTLs were identified for 890 proteins (26% of the measured proteome), an increase of 369 proteins over those reported by Sun *et al.* 2018^22^ using a genome-wide P-value threshold of P < 1.5×10^-11^. Significant *cis*-pQTLs mapped here are provided in **Supplementary Data 4**.

#### Mendelian randomisation

Genetic instruments for each aptamer were selected using a step-forward procedure that pruned the set of significant *cis*-pQTLs (**Supplementary Methods**) by linkage disequilibrium after mapping pQTL to GWAS summary statistics (**Supplementary Methods**). First, linkage disequilibrium was computed between all *cis*-pQTLs for each aptamer from the INTERVAL probabilistic genotype data using QCTOOL^139^ version 2 and LDstore^140^. For each aptamer and GWAS, *cis*-pQTLs were sorted by P-value and the variant with the smallest P-value added to the set of independent *cis*-pQTLs. Then, the remaining list of *cis*-pQTLs was filtered to remove any variants with r^2^ > 0.1 with any variants in the set of independent *cis*-pQTLs. This was repeated until no variants remained in the list of *cis*-pQTLs for the aptamer which mapped to the respective GWAS summary statistics (**Supplementary Table 6**).

Five Mendelian randomisation methods were used in parallel to estimate the causal effects of protein levels on disease (**Supplementary Table 5**), as each method makes different assumptions about instrument validity and are differentially robust to different sources of bias: (1) The inverse-variance weighted estimator, which takes the average causal estimate across all instruments: the average of their ratios of disease log odds over effect on protein levels multiplied by their standard errors^57^. It assumes all instruments are valid: the variant is associated with both the protein and disease, and that the variant has only a direct effect on that protein’s levels; it can only effect the disease by modifying the protein’s levels^57^. (2) The simple median estimator takes the median causal effect across instruments and is robust to outliers and up to 50% invalid instruments^58^. (3) The weighted median estimator, which weights each instrument by its inverse variance when calculating the median^58^. (4) The weighted mode estimator, which estimates the average causal effect from the largest set of instruments with consistent causal estimates^59^. (5) The Egger estimator which detects and corrects for bias arising from horizontal pleiotropy; where instruments are affecting other risk factors that may explain their observed association with the disease^60^. Where there were multiple aptamers meeting the criteria for Mendelian randomisation (**Online Methods**) for a given protein, causal estimates from the five Mendelian randomisation methods as well as the Egger intercept were averaged across the multiple aptamers (**Supplementary Table 5**, see also **Supplementary Methods** section above on averaging summary statistics across aptamers). A consensus causal estimate was subsequently obtained by taking the median estimate, standard error, 95% confidence interval, and P-value across the five Mendelian randomisation methods.

#### Colocalisation analysis

Bayesian colocalisation tests were performed between pQTL and GWAS signals to test whether both signals arose from a shared causal variant^62^. Tests were performed for *cis*-pQTLs used in Mendelian randomisation (**Supplementary Table 6**) with P-value < 1×10^-6^ in the respective GWAS, the suggested threshold for colocalisation testing^62^. For each test, SNPs within 200Kb of the *cis*-pQTL that mapped to the GWAS summary statistics (see above) were used. Colocalisation was tested using the R package coloc^62^ using summary-statistics based method (coloc.abf). For each test, five posterior probabilities were computed (**Supplementary Table 6**): PP.H0: no association with either the *cis*-pQTL or GWAS trait, PP.H1: significant *cis*-pQTL association but no association in GWAS, PP.H2: significant GWAS signal, but no association with the protein, PP.H3: *cis*-pQTL and GWAS signals arise from two independent causal variants, and PP.H4: *cis*-pQTL and GWAS signals arise from a shared causal variant. Evidence of colocalisation was defined as previously suggested^141^ as PP.H3 + PP.H4 ≥ 0.99 and PP4/PP3 ≥ 5: *i.e.* posterior probability of a shared causal variant 5-fold greater than posterior probability for associations arising from independent causal variants, and combined both posterior probabilities summing to 99%).

#### Drug target effect classification

Effects of drugs on PGS-associated proteins were classified into four categories: positive, negative, downstream, and unknown. Positive effects were those in which the drug increased the levels or activity of the protein: where the drug interaction was listed as an agonist, potentiator, inducer, or ligand of the protein. Negative effects were those in which the drug decreased the levels or activity of the protein: where the drug interaction was listed as an antagonist, inhibitor, or inhibitory allosteric modulator of the protein. Downstream effects were those in which different levels of the protein could affect drug efficacy: those where the protein was listed as a binder or carrier of the drug, where the drug was metabolised by the protein (enzyme substrate), or where the listed compound was also biomolecule a produced by the protein. The effect was considered unknown where the drug to protein interaction was listed as unknown/other, modulator, where an interaction was noted but no details on action were listed in the Drug Bank database^30^ version 5.17 released on the 2^nd^ of July 2020 (<https://go.drugbank.com/releases/latest>). Manual classification was performed based on available literature on drug to target interactions listed on DrugBank where the drug was listed as an enzyme substrate or target product. Drugs listed in **Table 1** were those with approved but not withdrawn status that (1) had a positive effect on the protein where decreased protein levels were associated with increased PGS, or (2) had a negative effect on the protein where increased protein levels were associated with increased PGS.

### Supplementary Tables

**Supplementary Table 1: Prevalent cardiometabolic disease events**

Hospital records in the 13.4 years prior to baseline assessment were summarised into 301 CALIBER phenotypes. Among those, we considered 48 to potentially indicate the presence of overt or sub-clinical cardiometabolic disease at baseline assessment and with potential to confound causal inference (**Fig. 2a**). Participants with hospital record for any of these events in the 13.4 years prior to baseline were excluded from analyses (**Online Methods**, N=87 of 3,174 participants excluded). This table details the 48 prevalent cardiometabolic disease phenotypes, the ICD-10 codes used to define them, and the number of participants hospitalised with that phenotype prior to baseline. Percentages given are relative to the total number of excluded participants. ICD-10 codes were obtained from https://github.com/spiros/chronological-map-phenotypes (git commit 32d55e48 on 23^rd^ March 2020).

Supplementary Table 2: Sensitivity and specificity of protein aptamers

For each PGS-associated protein, provides details of the intended target for each SomaLogic aptamer passing quality control (**Online Methods**). Aptamer: Sequence ID for the SomaLogic aptamer(s) targeting the protein. Intended Target: full name of the intended target for each aptamer as provided by SomaLogic. Note that these may be protein complexes or specific protein isoforms. UniProt: The UniProtKB identifier for the intended target protein. Entrez ID: the Entrez identifier in NCBI Gene. Chr: the chromosome the gene is located on, as listed on the NCBI gene page for the gene. Start: the start position of the gene on that chromosome on human genome the build GRCh37. Cross-reactivity: summary of cross-reactivity testing performed by SomaLogic for the aptamer against any proteins with >40% sequence homology. Entries may be a blank cell, indicating the aptamer has not yet been tested by SomaLogic, "no closely related human proteins" where the aptamer has been tested but there were no proteins with >40% sequence homology to the target protein, "no binding observed", "binding observed with at least 10x weaker affinity", or "Binding observed with similar affinity". Mass Spec confirmation: indicates whether binding of the aptamer to the target protein has been confirmed by mass spectrometry pulldown experiments either by SomaLogic (see the technical note published by SomaLogic at https://somalogic.com/technology/our‑platform/somamer‑specificity/) or by Emilsson *et al.* 2018^142^ (see Supplementary Tables 3 and 4 in Emilsson *et al.* 2018). cis pQTL: "yes" where aptamer had a significant *cis-*pQTL after hierarchical multiple testing correction (**Supplementary Information**), supporting the aptamer to target binding. In cases where aptamers bound another protein or differentially bound to specific protein isoforms the protein target in the main text (**Extended Data Table 3**) reflects this.

**Supplementary Table 3: Point estimates where multiple aptamers target a protein**

Details point estimates for PGS to protein to disease associations for the five PGS-associated proteins targeted by more than one aptamer. Columns are as described in **Extended Data Table 2** and **Extended Data Fig. 2**. Beta: standard deviation change in aptamer levels per standard deviation increase in PGS. 95% CI: 95% confidence interval. HR: hazard ratio for incident disease conferred per standard deviation increase in aptamer levels. OR: odds ratio for incident disease conferred through the protein per standard deviation increase in PGS. % PGS: Percentage of total effect of PGS on incident disease conferred through protein levels. The total odds ratio for myocardial infarction conferred per standard deviation increase in CAD PGS was 2.51 (95% CI: 1.50­­–4.22, P-value: 5×10^−4^). The total odds ratio for diabetes conferred per standard deviation increase T2D PGS was 1.97 (95% CI: 1.34­­–2.91, P-value: 6×10^−4^). Lighter text indicates P > 0.05 for the respective association. Under the “Effect of T2D PGS through BMI” heading, gives from mediation analysis (**Online Methods**), the estimated effect of T2D PGS on aptamer levels for WFIKKN2 through BMI (standard deviation change in aptamer levels through BMI per standard deviation increase of T2D PGS). Under the “Effect of T2D PGS independent of BMI” heading, gives from mediation analysis, the estimated effect of PGS for type 2 diabetes on protein levels independent of BMI. Lighter text indicates P > 0.05 for the respective association.

**Supplementary Table 4: Multivariable contributions of PGS and pQTLs to protein levels**

For each PGS associated protein details the independent contribution of PGS and pQTLs to protein levels. Joint beta estimates, their standard errors, 95% confidence intervals, and P-values are given for linear regression models fit for the indicated aptamer using the corresponding PGS and pQTLs as dependent variables. Coefficients are detailed for each aptamer separately, as lead pQTLs at each locus sometimes differed between aptamers targeting the same protein (**Supplementary Information**). Rows corresponding to pQTLs give their rsID in the "coefficient" column, its chromosome (chr) and position (pos) on build GRCh37, its effect allele (EA) whose dosage the beta estimate was fit for, the other allele (OA), its effect allele frequency (EAF), and whether the pQTL was a cis or trans-pQTL. Rows labelled "PGS" in the coefficient column give the effect of the corresponding PGS on the aptamer levels adjusting for the pQTLs. Beta coefficients correspond to standard deviation increase in aptamer levels per standard deviation increase in PGS levels, adjusting for the pQTLs, or standard deviation increase in aptamer levels per copy of the named effect allele adjusting for the PGS and other pQTLs.

Supplementary Table 5: Causal estimates from Mendelian randomisation analysis

From right to left: entries under the “Aptamer specific estimates” detail the causal estimates from five Mendelian randomisation methods for each PGS-associated protein aptamer with at least three independent genetic instruments (**Online Methods**, **Supplementary Table 6**), which were optimised per-aptamer (**Supplementary Information**). The (Intercept) entry indicates the estimate of the intercept in Egger regression. Under the “Causal estimate from each Mendelian randomisation method” heading, causal estimates for each method have been averaged where there were multiple aptamers targeting the protein (**Supplementary Information**). Entries under the “Consensus Causal Estimate” heading give the median estimated causal effect across the five Mendelian randomisation methods (excluding the Egger intercept term), also given in **Extended Data Fig. 5a**. The Pleiotropy P-value column corresponds to the P-value from the Egger intercept term, which indicates where P < 0.05, confounding of the causal estimate by associations between genetic instruments (*cis-*pQTLs) with multiple disease risk factors (horizontal pleiotropy). The Colocalisation column indicates whether there was evidence of pQTL and GWAS signals arising from a shared causal variant for any of the genetic instruments used for Mendelian randomisation (**Supplementary Information**, **Supplementary Table 6**). Under each heading, the "Estimate" column corresponds to the estimated causal effect on the log odds for disease risk per standard deviation increase in the protein/aptamer levels, and L95 and U95 the 95% confidence interval.

Supplementary Table 6: Genetic instruments for Mendelian randomisation analysis

Genetic instruments used in Mendelian randomisation analysis for aptamers with three or more independent by LD (r^2^ < 0.1) cis-pQTLs that mapped to disease GWAS summary statistics (**Supplementary Information**). Entries under the “Protein information” column give the chromosome and start location of the encoding gene on human genome build GRCh37. See **Extended Data Table 3** for further information on each protein. Entries under the "Aptamer QTL information" heading give details on the instruments used for the Mendelian randomisation tests for causal effects on the disease outcome listed under the Disease column. The chr and pos columns give the chromosome and position of the *cis*-pQTL. EA denotes the effect allele (minor allele in INTERVAL), OA the other allele, and EAF the effect allele frequency. The Beta column gives the standard deviation change in aptamer levels per copy of the effect allele in the pQTL summary statistics published by Sun *et al*. 2018. The SE column gives the standard error of the Beta estimate. Columns under the "GWAS QTL information" heading give details on the association of the instrument with log odds (log OR) of the respective disease outcome listed under the Disease column. The EAF column gives the frequency of the effect allele in the GWAS summary statistics. GWAS summary statistics were obtained from Nelson *et al.* 2017 for CAD (GCST004787), Wuttke *et al.* 2019 for CKD (GCST008065), Malik *et al.* in 2018 for IS (GCST006906) and Mahajan et al. 2018 for T2D (GCST007518). In all cases, we used the GWAS summary statistics for the samples of recent European ancestry. For T2D, we used the BMI-adjusted GWAS summary statistics in order to avoid false positive causal estimates arising where pQTL variants influence T2D risk through BMI rather than through the tested protein (horizontal pleiotropy). Colocalization of pQTL and GWAS signals was tested where the P-value < 1×10^-6^ for the instrument in the GWAS summary statistics (**Supplementary Information**). Colocalisation was tested using variants within 200Kb of the listed instrument. "Variants" lists the number of variants in that window that mapped between the GWAS and pQTL summary statistics. PP.H0, PP.H1, PP.H2, PP.H3, and PP.H4 give posterior probabilities from the colocalisation test for: (PP.H0) no association with either the cis-pQTL or GWAS trait, (PP.H1) significant cis-pQTL association but no association in GWAS, (PP.H2): significant GWAS signal, but no association with the protein, (PP.H3) cis-pQTL and GWAS signals arise from two independent causal variants, and (PP.H4) cis-pQTL and GWAS signals arise from a shared causal variant. Evidence of colocalisation (Yes/No in the colocalises column) was defined as PP.H3 + PP.H4 ≥ 0.99 and PP4/PP3 ≥ 5 (posterior probability of a shared causal variant 5-fold greater than posterior probability for associations arising from independent causal variants, and combined both posterior probabilities summing to 99%).

Supplementary Table 7: Drugs interacting with PGS-associated proteins

Information on the 236 drugs in DrugBank (https://go.drugbank.com/) that interact with PGS-associated proteins. Columns under the “Effect of drug on PGS-associated protein” heading detail the effect the listed drug has on the PGS-associated protein. Proteins may either be targets of the drug, carriers of the drug, enzymes modulated by the drug, or cellular transporters affected by the drug. Effects of drugs on PGS associated proteins (“Effect direction”) were classified into four categories: positive, negative, downstream, and unknown. Positive where the drug increased the levels or activity of the protein, negative where the drug decreased the levels or activity of the protein, downstream where the protein levels effect the drug rather than the drug effecting the protein levels, and unknown where the direction could not be determine (**Supplementary Information**). The “Drug effect opposite to PGS” column indicates whether the direction of effect of the drug on protein levels is inverse to the effect of PGS on protein levels. The “Method of drug action” column indicates whether the PGS-associated protein is among the proteins through which the drug’s method of pharmacological action is known to take place. Columns under the “Drug details” heading give the description of the drug provided by DrugBank, the type of the drug (small molecule or biotech), and the groups the drug belongs to in DrugBank (approved, nutraceutical, investigational, experimental, and/or withdrawn). Columns under the “Pharmacology” heading provided information from the following fields in DrugBank. “Indication”: a descriptive field describing the approved conditions, diseases, or states for which a drug can safely and effectively be used. “Pharmacodynamics”: A description of how the drug modifies or affects the body. “Mechanism”: a description of the biochemical interaction through which the drug produces its intended effect. Columns under the “Proteins interacting with drug” heading provide a listing of human proteins that are known to interact with the drug in some way. Columns and rows may need to be resized to view the full contents of descriptive fields of interest. Pharmacological targets: a comma separate list of drug to molecule (e.g. protein) binding or interactions describing how the drug exerts its pharmacological action. Other targets: a comma separate list of drug to molecule (e.g. protein) binding or interactions excluding targets through which the drug exerts its pharmacological action (targets to which the drug binds but has no effect, or unknown effect). Enzymes: a comma separated list of enzymes (e.g. proteins) which metabolise the drug. Carriers: a list of molecules (e.g. proteins) that bind the drug and facilitate its movement within the body to the target molecule. Transporters: a comma separated list of cell membrane bound proteins that facilitate transport of the drug into and out of cells. Columns under the “Maximum phase clinical trials” heading provide details on the set of clinical trials that have reached (or are underway) the most advanced phase among all clinical trials listed on the DrugBank page for the drug. Max phase: maximum phase reached in any clinical trial listed under the Clinical Trials heading on the drugs DrugBank page. Status: status or outcome of that clinical trial. Purpose: purpose of that clinical trial. Indications: list of conditions the clinical trial was testing the drug for. Distinct clinical trials are separated by a “ | “, while clinical trials that target multiple conditions have those conditions separated by a “ / “. Finally, the “Clinical trials relevant to PGS associated with protein interacting with drug” column provides details on clinical trials of any phase that have been undertaken for conditions relating to the listed PGS on that row.

### Supplementary Data

Supplementary Data 1: Curated SomaLogic aptamer information.

Annotations for the 4,034 SOMAscan aptamers quantified in the INTERVAL cohort. QC tag: reason for aptamer exclusion. The 3,793 high-quality aptamers (**Online Methods**) are tagged “Protein”, with other values indicating the exclusion reason provided by SomaLogic. Under the “Protein Information” heading: full name of the intended target for each aptamer as provided by SomaLogic. Note that these may be protein complexes or specific protein isoforms. UniProt: The UniProtKB identifier for the intended target protein. Under the “Encoding Gene” heading: Gene: Gene(s) encoding the targeted protein. Entrez: the Entrez identifier in NCBI Gene. Chr: the chromosome the gene is located on, as listed on the NCBI gene page for the gene. Start: the start position of the gene on that chromosome on human genome the build GRCh37. Where multiple proteins were targeted, their corresponding UniProt identifiers and encoding genes are separated by a "|", as are their respective chromosomes, start, and end positions on build GRCh37. Genes with multiple start sites have these separated by a ";". Under the “aptamer specificity” heading: Cross-reactivity: summary of cross-reactivity testing performed by SomaLogic for the aptamer against any proteins with >40% sequence homology. Entries may be a blank cell, indicating the aptamer has not yet been tested by SomaLogic; “under investigation”, indicating ongoing experiments by SomaLogic; "no closely related human proteins" where the aptamer has been tested but there were no proteins with >40% sequence homology to the target protein; "no binding observed", "binding observed with at least 10x weaker affinity", or "Binding observed with similar affinity" for proteins with >40% sequence homology with the intended target. Mass Spec confirmation: indicates whether binding of the aptamer to the target protein has been confirmed by mass spectrometry pulldown experiments either by SomaLogic (see the technical note published by SomaLogic at https://somalogic.com/technology/our‑platform/somamer‑specificity/) or by Emilsson *et al.* 2018 (see Supplementary Tables 3 and 4 in Emilsson *et al.* 2018). cis pQTL: "yes" where aptamer had a significant *cis-*pQTL after hierarchical multiple testing correction (**Supplementary Methods**; **Supplementary Data 4**), supporting the aptamer to target binding. Under the “INTERVAL cohort variable mapping” heading: “Column name”: column name in the post-QC dataset generated by Sun *et al.* 2018 (see **Data Availability**). “pQTL summary statistics”: name of the aptamer in the pQTL summary statistics published by Sun *et al.* 2018 (see **Data Availability**).

Supplementary Data 2: Aptamer distributions and covariate associations

**a)** Aptamer distributions and covariate associations in the raw data. For each aptamer, summarises the distribution of the aptamer's levels prior to quality control, covariate adjustment, and normalisation (columns F-M), and details the association between the raw aptamer levels and the covariates the aptamer levels were adjusted for prior to inverse rank normalisation by Sun *et al.* 2018 (age, sex, first three genotype PCs, time between blood draw and sample processing >1 day, columns N-AK), sample measurement batch (columns BN-BQ) which were regressed out here (**Online Methods**) from the inverse rank normalised data produced by Sun *et al.* 2018, and covariates adjusted for in sensitivity analysis (**Extended Data Fig. 2f**) (BMI, season, time of day; columns BR-ES). The Aptamer Annotation columns (columns B-D) provide details on the intended target of each Aptamer (column A): the intended protein target as provided by SomaLogic, the UniProt KB identifier for the target protein(s), and the gene(s) encoding the protein. Where multiple proteins were targeted, their corresponding UniProt identifiers and encoding genes are separated by a "|", as are their respective chromosomes and start positions on build GRCh37. Genes with multiple start sites have these separated by a ";". Under the "Summary of apatamer level distribution" heading: Minimum: the smallest value for the aptamer level in the 3,087 samples without prevelant cardiometabolic disease. N < 5 SD: number of samples at least 5 standard deviations below the mean. N > 5 SD number of samples at least 5 standard deviations above the mean. Under the covariate association headings: Beta: beta estimate in linear regression model fit for the apatamer levels on that covariate indicating change in protein levels per unit increase in the covariate. L95: lower 95% confidence interval. U95: upper 95% confidence interval. P-value: p-value for the beta-estimate in linear regression. Point estimates are shown in black where P < 0.05, and are dulled in grey where P > 0.05. For covariates that were fit as factors (sex, batch, time between blood draw and sample processing, date of sample collection, time of day of sample collection) the group to which the beta estimate pertains is detailed in the second header row, as is the reference group to which it is compared to in the model. Aptamers pertaining to the PGS associated proteins (**Supplementary Table 2**) are detailed at the top of the table, and are indicated by the “Associated with PGS” column (column E). Proteins are ordered top to bottom in ascending order by association with any PGS (smallest FDR correct P-value across the five PGSs from **Supplementary Data 3a**).

**b)** Aptamer distributions and covariate associations after quality control and normalisation. Columns are as described in **Supplementary Data 2a**. Here, the unit of effects of covariates on normalised aptamer levels (Beta, 95% confidence interval) are standard deviation change in protein levels per unit increase in covariates.

Supplementary Data 3: Summary statistics for statistical tests.

Each sheet **a-h** contains a distinct set of summary statistics. The figures which they underlie are noted in the first sentence of each caption.

**a)** Summary statistics underlying **Fig. 1a**. Details associations between the each of the five cardiometabolic PGSs with each of the 3,793 SomaLogic aptamers targeting 3,438 proteins. Under the “Protein Information” heading: full name of the intended target for each aptamer as provided by SomaLogic. Note that these may be protein complexes or specific protein isoforms. UniProt: The UniProtKB identifier for the intended target protein. Entrez ID: the Entrez identifier in NCBI Gene. Chr: the chromosome the gene is located on, as listed on the NCBI gene page for the gene. Start: the start position of the gene on that chromosome on human genome the build GRCh37. Where multiple proteins were targeted, their corresponding UniProt identifiers and encoding genes are separated by a "|", as are their respective chromosomes and start positions on build GRCh37. Genes with multiple start sites have these separated by a ";". Under the “Protein association” heading: point estimates of the effect of the PGS on the given aptamer target. Point estimates were averaged (**Supplementary Methods**) where multiple high-quality aptamers target the protein (with point estimates duplicated across rows for the N > 1 aptamers). Beta: standard deviation change in protein levels per standard deviation increase in PGS. L95 and U95 give the lower and upper bounds of the 95% confidence interval. FDR: false discovery rate adjusted P-value (adjusted across all 3,438 proteins for each PGS separately).

**b)** Summary statistics underlying **Extended Data Fig. 2a-c.** Details PGS to protein associations assessed using orthogonal proteomics technology (Olink Explore and Olink T96). Note each figure panel comprises PGS to protein associations assessed in different groups of samples using different proteomics technology, see **Extended Data Fig. 2** for details. Point estimates (Beta) indicate the standard deviation change in protein levels per standard deviation increase in the PGS when assessed using the orthogonal proteomics technology.

**c)** Summary statistics underlying **Extended Data Fig. 2f.** Details PGS to protein associations adjusting for (1) circadian effects, (2) seasonal effects, (3) when including participants with prevalent cardiometabolic disease, and (4) adjusting for BMI. Under the “Protein association” heading: point estimates of the effect of the PGS on the given aptamer target. Point estimates were averaged (**Supplementary Methods**) where multiple high-quality aptamers target the protein (with point estimates duplicated across rows for the N > 1 aptamers). Beta: standard deviation change in protein levels per standard deviation increase in PGS in the given sensitivity analysis. L95 and U95 give the lower and upper bounds of the 95% confidence interval. Under the “Aptamer association” heading: point estimates for the effect of PGS on specific aptamers for the given target. Identical to information under the “Protein association” heading where the protein is targeted by one aptamer. Aptamer: Sequence ID for the SomaLogic aptamer(s) targeting the protein.

**d)** Summary statistics underlying **Extended Data Fig. 2g.** Extended details from mediation analysis of T2D PGS to protein associations by BMI. The natural direct effect corresponds to the effect of T2D PGS on protein levels independent of BMI. The natural indirect effect corresponds to the effect of T2D PGS on protein levels through BMI. The total effect, not detailed in **Extended Data Fig. 2g,** corresponds to the sum of the natural direct effect and natural indirect effect. Units of point estimates are standard deviation change in protein levels per standard deviation increase in T2D PGS. The % of total effect was calculated by dividing the corresponding point estimate by the point estimate for the total effect for the PGS to protein association. Point estimates under the “Aptamer association” heading are identical to those under the “Protein association” heading where only one aptamer targeted the protein (all proteins except WFIKKN2).

**e)** Summary statistics underlying **Extended Data Fig. 3** and polygenicity calculations. Each PGS was partitioned into 1,703 approximately independent LD blocks (**Online Methods**) then tested for association with each protein. Columns under the “LD block” heading provide information on the genomic location (human genome build GRCh37) and total size (in base pairs) for each of these 1,703 LD blocks. Columns under the “Protein association” detail associations between the protein levels and the subset of the PGS computed using only variants from this LD block. Units of point estimates are standard deviation change in protein levels per standard deviation increase in PGS LD block. Point estimates under the “Aptamer association” heading give the association for each individual aptamer targeting the listed protein, and are identical to those under the “Protein association” heading where only one aptamer targeted the protein.

**f)** Summary statistics underlying polygenicity calculations shown in **Fig 1c.** Following on from **Supplementary Data 3e**, LD blocks were removed from each PGS to protein association in ascending order of P-value (see **Supplementary Data 3e**) until the PGS to protein association was attenuated (P-value > 0.05). This table details the PGS to protein association at each step in this process until the association was attenuated. At each step, the number of LD blocks removed from the PGS are given, along with the total base pairs in the genome these LD blocks covered, and the total % of the genome removed from the PGS. Point estimates under the “Aptamer association” are identical to those under the “Protein association” heading where only one aptamer targeted the protein. The final row for each PGS to protein association, where the P-value > 0.05, gives the polygenicity of the PGS to protein association in the % removed column.

**g)** Summary statistics underlying **Fig. 2b**. Associations between protein levels and incident MI and incident T2D over 7.7 years of follow-up. HR: hazard ratio for incident disease adjusting for age and sex conferred per standard deviation increase in the protein/aptamer levels.

**h)** Summary statistics underlying **Fig. 2d**. Point estimates from mediation analysis of PGS to incident disease associations by PGS-associated proteins. The natural direct effect corresponds to the effect of the PGS on incident disease independent of the given protein. The natural indirect effect corresponds to the effect of PGS on incident disease through the given protein. The total effect corresponds to the sum of the natural direct effect and natural indirect effect. Units of point estimates are odds ratio (OR) for disease adjusting for age and sex per standard deviation increase of the PGS. The % of total effect was calculated by dividing the corresponding point estimate by the point estimate for the total effect for the PGS to incident disease association.

Supplementary Data 4: Hierarchically significant cis-pQTLs.

For each aptamer, pQTLs have been filtered to bi-allelic SNPs, then pruned by linkage disequilibrium (LD) so that all pQTLs for each aptamer have r^2^ < 0.1 with each other. A/T and G/C SNPs were also removed prior to LD pruning where there minor allele frequency was > 42%. Under the “Protein Information” heading: full name of the intended target for each aptamer as provided by SomaLogic. Note that these may be protein complexes or specific protein isoforms. UniProt: The UniProtKB identifier for the intended target protein. Entrez ID: the Entrez identifier in NCBI Gene. Chr: the chromosome the gene is located on, as listed on the NCBI gene page for the gene. Start: the start position of the gene on that chromosome on human genome the build GRCh37. Where multiple proteins were targeted, their corresponding UniProt identifiers and encoding genes are separated by a "|", as are their respective chromosomes and start positions on build GRCh37. Genes with multiple start sites have these separated by a ";". For each Aptamer, *cis-*pQTLs were mapped within 1Mb of the listed start positions and chromosomes for the Gene(s) coding for the target protein. Columns under the "pQTL Information" heading provided details on the pQTLs, including their chromosome (Chr), position on build GRCh37 (Pos) effect allele (EA, always minor allele), other allele (OA), frequency of the effect allele (EAF), effect size (Beta) corresponding to standard deviation change in aptamer levels per copy of the effect allele, standard error of the effect size (SE), P-value, the locally corrected P-value in the hierarchical correction (P-value adjusted for the number of variants within 1Mb of any start site for any gene coding for the target protein), and the globally corrected P-value for the aptamer in hierarchical correction (smallest P-value per aptamer FDR corrected across all aptamers; **Supplementary Methods**). Beta, SE, and P-value were obtained from summary statistics published by Sun *et al*. 2018.
